# Adventures in Multi-Omics I: Combining heterogeneous datasets via relationships matrices

**DOI:** 10.1101/857425

**Authors:** Deniz Akdemir, Ron Knox, Julio Isidro-Sánchez

## Abstract

Private and public breeding programs, as well as companies and universities, have developed different genomics technologies which have resulted in the generation of unprecedented amounts of sequence data, which bring new challenges in terms of data management, query, and analysis. The magnitude and complexity of these datasets bring new challenges but also an opportunity to use the data available as a whole. Detailed phenotype data, combined with increasing amounts of genomic data, have an enormous potential to accelerate the identification of key traits to improve our understanding of quantitative genetics. Data harmonization enables cross-national and international comparative research, facilitating the extraction of new scientific knowledge. In this paper, we address the complex issue of combining high dimensional and unbalanced omics data. More specifically, we propose a covariance-based method for combining partial datasets in the genotype to phenotype spectrum. This method can be used to combine partially overlapping relationship/covariance matrices. Here, we show with applications that our approach might be advantageous to feature imputation based approaches; we demonstrate how this method can be used in genomic prediction using heterogenous marker data and also how to combine the data from multiple phenotypic experiments to make inferences about previously unobserved trait relationships. Our results demonstrate that it is possible to harmonize datasets to improve available information across gene-banks, data repositories or other data resources.

**Key message:** Several covariance matrices obtained from independent experiments can be combined as long as these matrices are partially overlapping. We demonstrate the usefulness of this methodology with applications in combining data from several partially linked genotypic and phenotypic experiments.

**Author contribution statement:** –DA: Conception or design of the work, statistics, R programs, simulations, drafting the article, and critical revision of the article.

–JIS: R programs, graphs, drafting the article, critical revision of the article.

–RK: Critical revision of the article.

## 1 Introduction

The rapid scientific progress in these genomic approaches is due to the decrease in genotyping costs by the development of next generation sequencing platforms since 2007 [Mardis, 2008a,b]. High-throughput instruments are routinely used in laboratories in basic science applications, which has led to the democratization of genome-scale technologies, such as genomic predictions and genome-wide associating mapping studies. Genomic prediction, i.e. predicting an organism’s phenotype using genetic information [Meuwissen et al., 2001], it is currently used by many breeding companies, because it improves three out of the four factors affecting the breeder equation [Hill and Mackay, 2004]. It reduces generation number, improves accuracy of selection, and increases selection intensity for a fixed budget when comparing with marker-assisted selection or phenotypic selection [Desta and Ortiz, 2014, Heffner et al., 2011, 2010, Juliana et al., 2018, de los Campos et al., 2013]. Genomic prediction and selection (GS) are a continuously progressing tool that promises to help meet the human food challenges in the next decades [Crossa et al., 2017]. Genome-wide associating mapping studies, which originated in human genetics [Bodmer, 1986, Risch and Merikangas, 1996, Visscher et al., 2017], have also become a routine in plant breeding [Gondro et al., 2013].

The biological data generated in the last few years from this genomic progress have grown exponentially which have led to a high dimensional and unbalanced nature of the ‘omics’ data. Data normally comes in various forms of marker and sequence data: expression, metabolomics, microbiome, classical phenotype, image-based phenotype [Bersanelli et al., 2016]. Private and public breeding programs, as well as companies and universities, have developed different genomics technologies which have resulted in the generation of unprecedented levels of sequence data, which bring new challenges in terms of data management, query, and analysis.

It is clear that detailed phenotype data, combined with increasing amounts of genomic data, have an enormous potential to accelerate the identification of key traits to improve our understanding of quantitative genetics [Crossa et al., 2017]. Nevertheless, one of the challenges that still needs to be addressed is the incompleteness inherent in these data, i.e., several types of genomic/phenotypic information covering only a few of the genotypes under study [Berger et al., 2013]. Data harmonization enables cross-national and international comparative research, as well allows the investigation of whether or not datasets have similarities. In this paper, we address the complex issue of utilizing the high dimensional and unbalanced omics data by combining the relationship information from multiple data sources, and how we can facilitate data integration from interdisciplinary research. The increase of sample size and the improvement of generalizability and validity of research results constitute the most significant benefits of the harmonization process. The ability to effectively harmonize data from different studies and experiments facilitates the rapid extraction of new scientific knowledge.

One way to approach the incompleteness and the disconnection among datasets is to combine the relationship information learned from these datasets. The statistical problem addressed in this paper is the calculation of a combined covariance matrix from incomplete and partially overlapping pieces of covariance matrices that were obtained from independent experiments. We assume that the data is a random sample of partial covariance matrices from a Wishart distribution, then we derive the expectation-maximization algorithm for estimating the parameters of this distribution. According to our best knowledge no such statistical methodology exists, although the proposed method has been inspired by similar methods such as (conditional) iterative proportional fitting for the Gaussian distribution [Cramer, 1998, 2000] and a method for combining a pedigree relationship matrix and a genotypic matrix relationship matrix which includes a subset of genotypes from the pedigree-based matrix [Legarra et al., 2009] (namely, the H-matrix). The applications in this paper are chosen in the area of plant breeding and genetics. However, the statistical method is applicable much beyond the described applications in this article.

Integration of heterogeneous and large omics data constitutes a challenge and an increasing number of scientific studies address this issue. A brief review and classification of some promising statistical approaches are described in Bersanelli et al. [2016]. According to this article, our covariance based method falls in the network based data integration category (as opposed to non-network based methods such as feature imputation) which include popular methods such as similarity network fusion Wang et al. [2014], weighted multiplex networks Menichetti et al. [2014] both of which can be used to combine several **complete** networks by suitable weighting. The main breakthrough here is that the proposed method in this article can be used to combine several **incomplete but partially overlapping** networks and that the proposed approach is supported theoretically by the maximum likelihood formalization.

## 2 Methods and Materials

### 2.1 Statistical methods for combining incomplete data

#### 2.1.1 Imputation

The standard method of dealing with heterogeneous data involves the imputation of features [Shrive et al., 2006]. If the datasets to be combined overlap over a substantial number of features then the unobserved features in these datasets can be accurately imputed based on some imputation method [Bertsimas et al., 2017].

Imputation step can be done using many different methods: Several popular approaches include Beagle [Browning and Browning, 2016], random forest [Breiman, 2001] imputation, expectation maximization based imputation [Endelman, 2011], low-rank matrix factorization methods that are implemented in the R package [Hastie and Mazumder, 2015]. In addition, parental information can be used to improve imputation accuracies [Nicolazzi et al., 2013, Gonen et al., 2018, VanRaden et al., 2015, Browning and Browning, 2009]. In this study, we used the low-rank matrix factorization method in all of the applications which included an imputation step. The selection of this method was due to computational burden of the other alternatives.

#### 2.1.2 Combining genomic relationship matrices

In this section, we describe the Wishart EM-Algorithm for combining partial genetic relationship matrices^1^.

**Wishart EM-Algorithm for Estimation of a Combined Relationship Matrix from Partial Samples**

Let *A* = *{a*_1_, *a*_2_, …, *a*_*m*_*}* be the set of not necessarily disjoint subsets of genotypes covering a set of *K* (i.e., 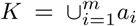) with total *n* genotypes. Let 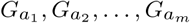 be the corresponding sample of genetic relationship matrices (kernels).

Starting from an initial estimate of the genetic relationship matrix *Σ*^(0)^ = *νΨ* ^(0)^, the Wishart EM-Algorithm repeats updating the estimate of the genetic relationship matrix until convergence:

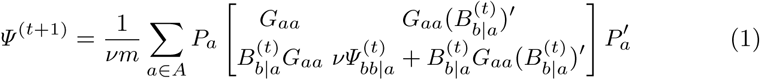

Where 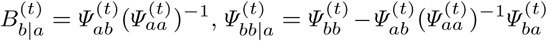, *a* is the set of genotypes in the given partial genomic relationship matrix and *b* is the set difference of *K* and *a*. The matrices *Pa* are permutation matrices that put each matrix in the sum in the same order. The initial value, *Σ*^(0)^ is usually assumed to be an identity matrix of dimesion *n*. The superscripts in parenthesis ‘(*t*)’ denote the iteration number. The estimate *Ψ* ^(*T*)^ at the last iteration converts to the estimated genomic relationship with *Σ*^(*T*)^ = *νΨ* ^(*T*)^.

A weighted version of this algorithm can be obtained replacing *Gaa* in Equation 1 with 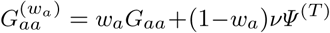 for a vector of weights (*w*_1_, *w*_2_, …, *w*_*m*_)*′*.

Derivation of the Wishart-EM algorithm and and its asymptotic errors are given in Supplementary.

### 2.2 Materials: datasets and experiments

In this section, we describe the datasets and the experiments we have designed to explore and exploit the Wishart EM-Algorithm.

Note that the applications in the main text involve real datasets and validation with such data can only be as good as the ground truth known about the underlying system. We also included several simulation studies in the Supplementary (Supplementary Applications 1 and 2) using simulated data to show that the algorithm performs as expected (maximizes the likelihood and provides a ‘good’ estimate of the parameter values) when the ground truth is known.

#### Application 1-Potato dataset; when imputation is not an option. Anchoring independent pedigree-based relationship matrices using a genotypic relation matrix

The Wishart EM-Algorithm can be used when the imputation of the original genomic features is not feasible. For instance, it can be used to combine partial pedigree-based relationship matrices with marker-based genomic relationship matrices. In this application, we demonstrate that genomic relationship matrices can be used to connect several pedigree-based relationship matrices.

The dataset is cited in [Endelman et al., 2018] and is available in the R Package AGHmatrix [Rampazo Amadeu et al., 2016]. It consists of the pedigree of 1138 potato genotypes, 571 of these genotypes also have data for 3895 tetraploid markers. The pedigree-based relationship matrix A was calculated with R package AGHmatrix [Rampazo Amadeu et al., 2016] using pedigree records, there were 185 founders (clones with no parent).

At each instance of the experiment, two non-overlapping pedigree-based relationship matrices each with the sample size *Nped ∈* {100, 150, 250} genotypes selected at random from the were 571 genotypes were generated. In addition, a genotypic relationship matrix was obtained for a random sample of *Ngeno ∈* {20, 40, 80} genotypes selected at random half from the genotypes in the first pedigree and a half from the genotypes from the second pedigree. These genetic relationship matrices were combined to get a combined genetic relationship matrix (See Figure 1). This combined relationship matrix was compared to the pedigree-based relationship matrix of the corresponding genotypes using mean squared errors and Pearson’s correlations. This experiment was repeated 30 times for each *Ngeno, Nped* pair.

**Fig. 1.**
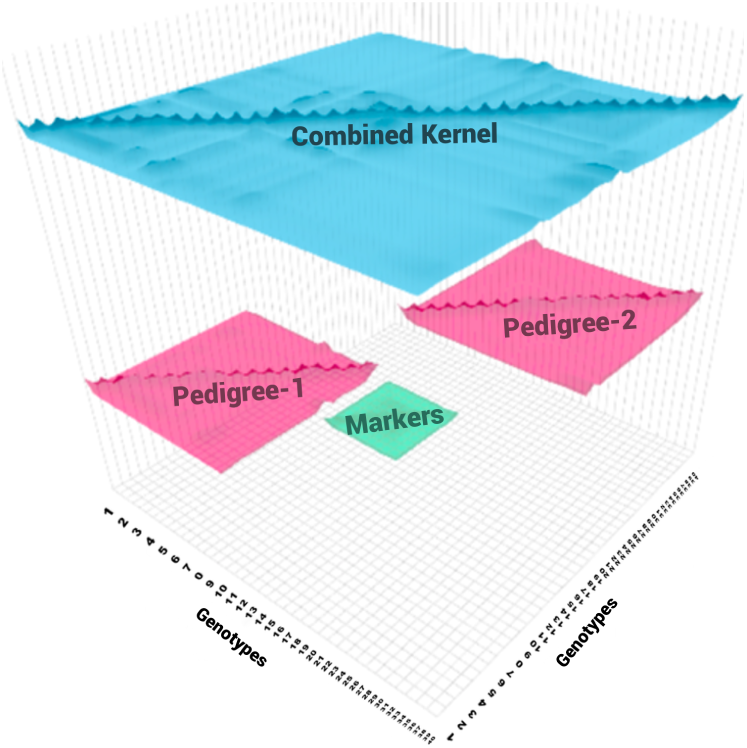
Application 1: At each replication of the experiment, in pink two non-overlapping pedigree-based relationship matrices are selected at random (20 individuals each) from the 571 genotypes. In green, a genomic relationship matrix obtained from a random sample of genotypes, half from the genotypes in the first pedigree (10) and half from the genotypes from the second pedigree (10). These three relationship matrices were combined to get a combined relationship matrix (in blue).

#### Application 2 - Rice dataset. Combining independent low density marker datasets

Rice dataset was downloaded from www.ricediversity.org. After curation, the marker dataset consisted of 1127 genotypes observed for 387161 markers. We treat the totality of information as the ground truth, i.e., we assume that the true genomic relationship for the 1127 genotypes is characterized by the 387161 markers. The purpose of this application is demonstrate that we can make inferences about the assumed true genomic relationship matrix by observing several smaller heterogeneous subsets of the available. This involves inferring a common estimate for the relationships that are already observed and producing estimates for relationships that haven’t been observed. Supplementary Figure S5 demonstrate this experiment pictorially.

In each instance of the experiment, *N*_*Kernel*_ *∈* {3, 5, 10, 20, 40, 80} marker datasets with 200 genotypes and 2000 markers were created by randomly sampling the genotypes and markers in each genotype file. These datasets were combined using the Wishart EM-Algorithm and also by imputation to give two genomic relationship matrices. For the totality of genotypes in these combined datasets, we also randomly sampled 2000, 5000 or 10000 markers and calculated the genomic relationships based on these marker subsets. All of these genomic relationship matrices were compared with the corresponding elements of the relationship matrix based on the entire genomic data by calculating the mean squared error between the upper diagonal elements including the diagonals. This experiment was replicated 20 times. Application results are showed in Figure 7.

#### Application 3 - Wheat Data at Triticeae Toolbox. Combining genomic datasets to use in genomic prediction

This application involves estimating breeding values for seven economically important traits for 9102 wheat lines obtained by combining 16 publicly available genotypic datasets. The genotypic and phenotypic data were downloaded from the Triticeae toolbox database. Each of the marker datasets were pre-processed to produce the corresponding genomic relationship matrices. Table 1 and Supplementary Figure S7 describes the phenotypic records and number of distinct genotypes for each trait.

**Table 1.**
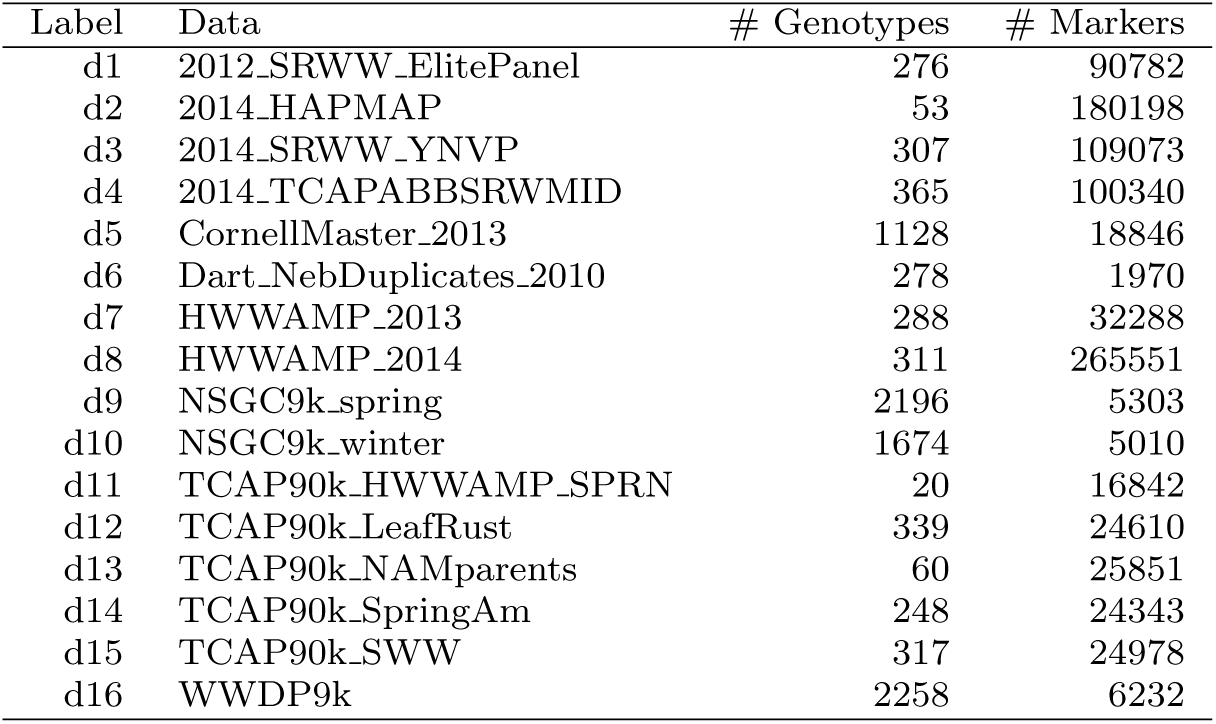
Marker datasets from Triticeae Toolbox: Labels and names for the datasets, number of genotypes and markers in each of the selected 16 genotypic datasets.

Using the combined relationship matrix we can build genomic prediction models. To test the performance of predictions based on the combined relationship matrix, we formulated two cross-validation scenarios. The intersection of genotypes among the 16 genotypic experiments is shown in Figure 2 and the intersection of common markers among genotypic experiments in Figure 3.

**Fig. 2.**
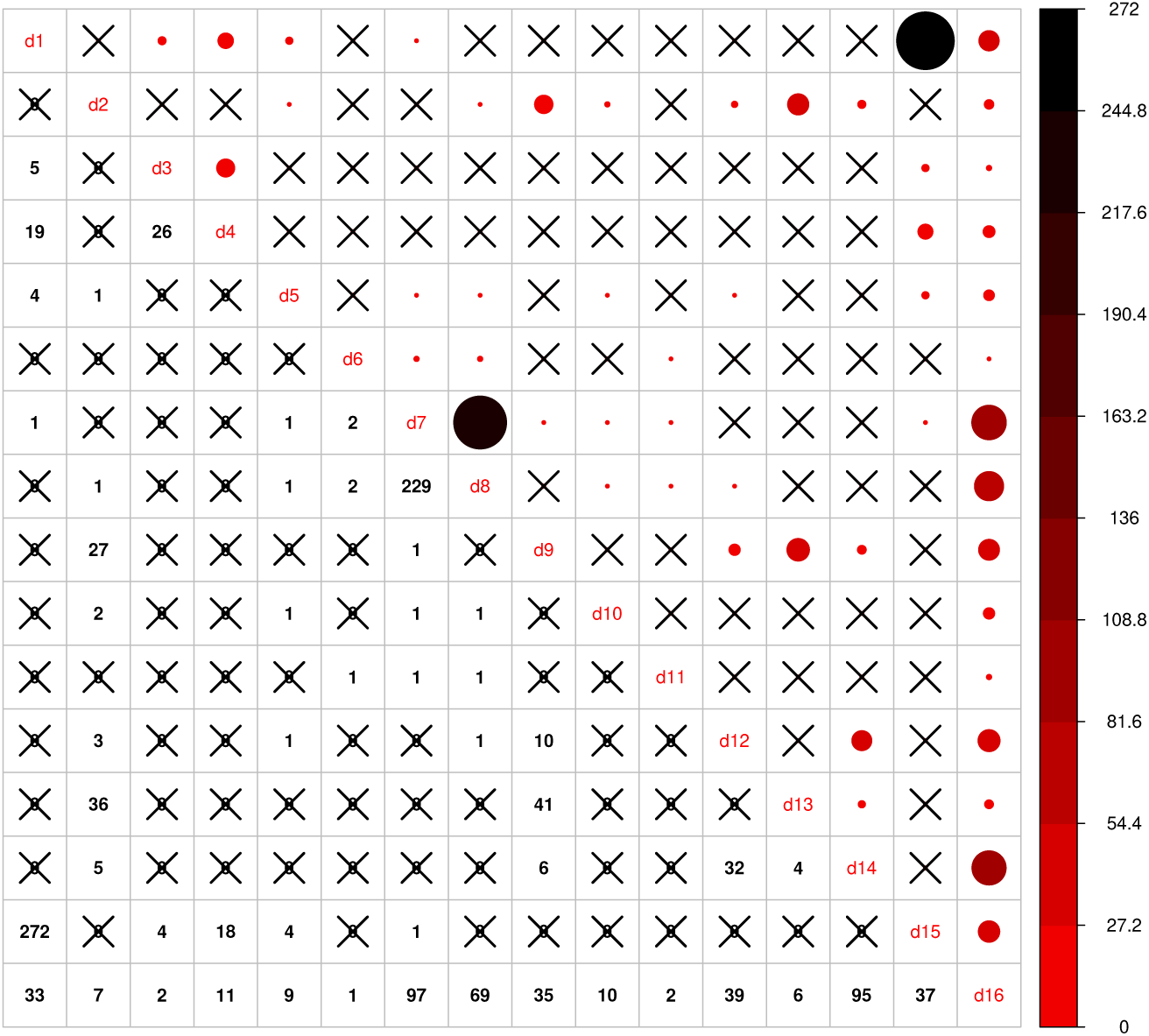
Intersection of genotypes among 16 genotypic experiments. The number of common genotypes among the 16 genotypic datasets are given on the lower diagonal, no intersection is marked by ‘X’. Upper diagonal of the figure gives a graphical representation of the same, larger circles represent higher number of intersections.

**Fig. 3.**
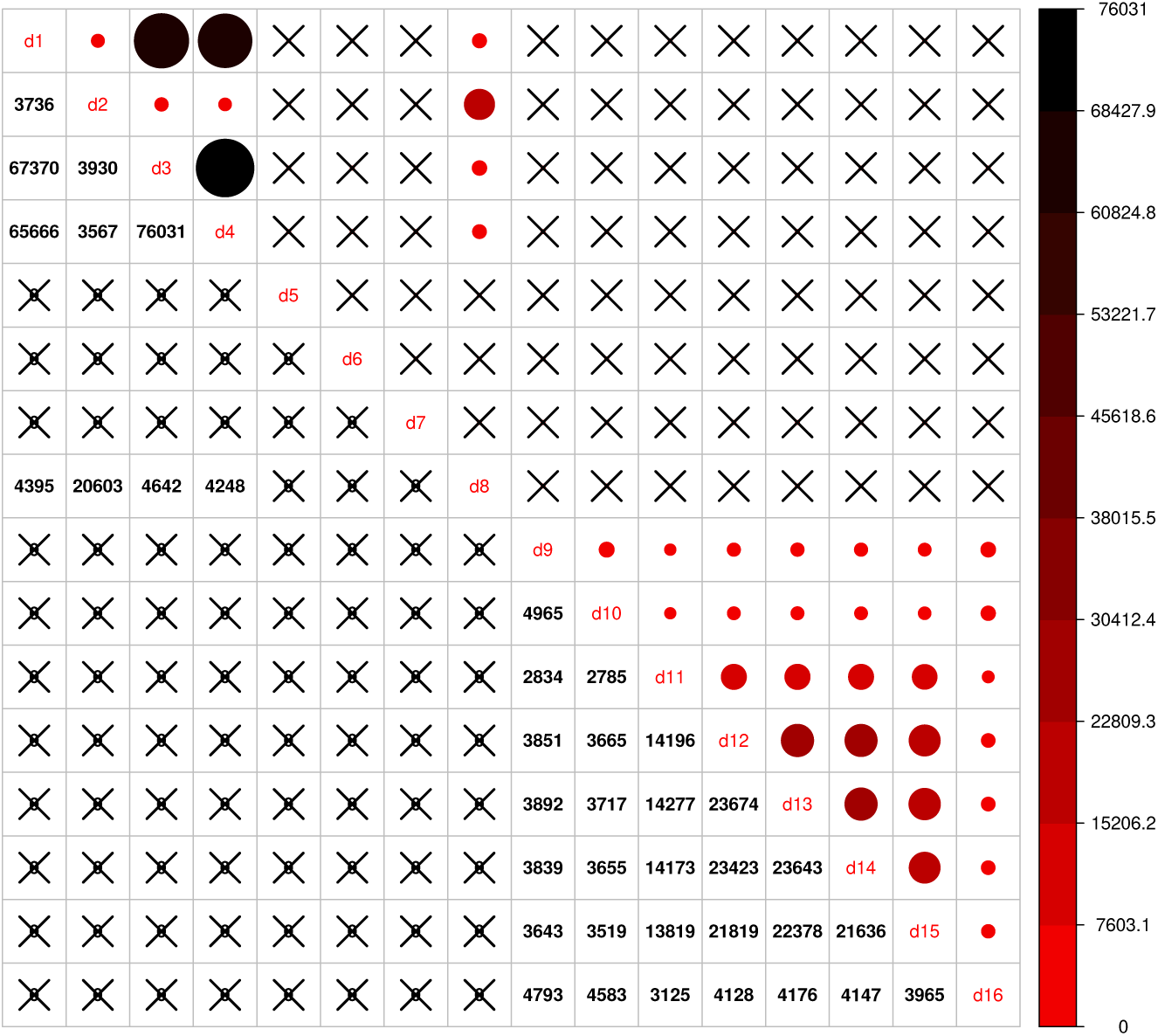
Intersection of markers among 16 genotypic experiments. The number of common markers among the 16 genotypic datasets are given on the lower diagonal, no intersection is marked by ‘X’. Upper diagonal of the figure gives a graphical representation of the same thing.

**Fig. 4.**
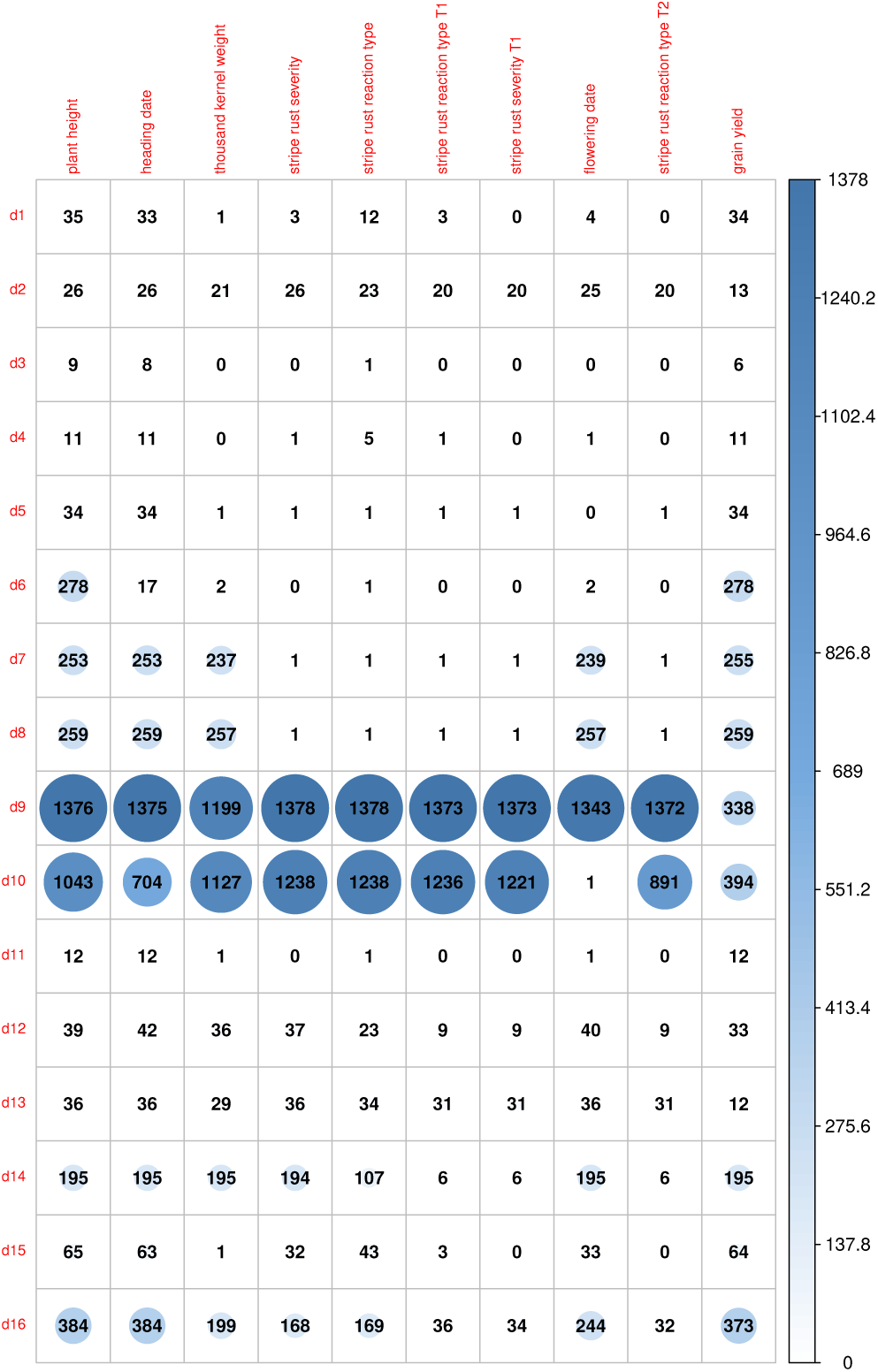
Availability of phenotypic data for the genotypes in 16 genotypic datasets for 10 traits. Here we indicated the traits with most phenotypic records for the genotypes in the 16 genotypic datasets.Plant height, grain yield, and heading time are the most measured trait across all the environments. Some trials have few measures. This graph shows the unbalanced and the need for harmonization of datasets.

##### Cross-validation scenario 1

The first scenario involved a 10 fold cross-validation based on a random split of the data. For each trait, the available genotypes were split into 10 random folds. The GEBVs for each fold were estimated from a mixed model (see Supplementary Section 5.4 for a description of this model) that was trained on the phenotypes available for the remaining genotypes. The accuracy of the predictions was evaluated by calculating the correlations between the GEBVs and the observed trait values.

##### Cross-validation scenario 2

Here, we performed a leave one dataset out cross-validation. i.e. we leave out the phenotypic records of the associated genotypes in one of the 16 genomic datasets and then estimate the trait values of those genotypes based on the basis of a mixed trained model. The training population was built on the remaining genotypes and phenotypic information after leaving the phenotypic records out. This scenario was used for each trait, and the accuracies were evaluated by calculating the correlations between the estimated and the observed trait values within each dataset.

#### Application 4 - Wheat Data at Triticeae Toolbox. Combining Phenotypic Experiments

The Wishart EM-Algorithm can also be used to combine correlation matrices^2^ obtained from independent phenotypic experiments. One-hundred forty four phenotypic experiments involving 95 traits in total were selected from 2084 trials and 216 traits available at the Triticeae Toolbox. In this filtered set of trials, each trial and trait combination had at least 100 observations and two traits. Furthermore, the percentage of missingness in these datasets was at most 70%. The mean and the median of the number of traits in these trials were 5.9 and 4 correspondingly (See Figure 5 and Supplementary Figure S6).

**Fig. 5.**
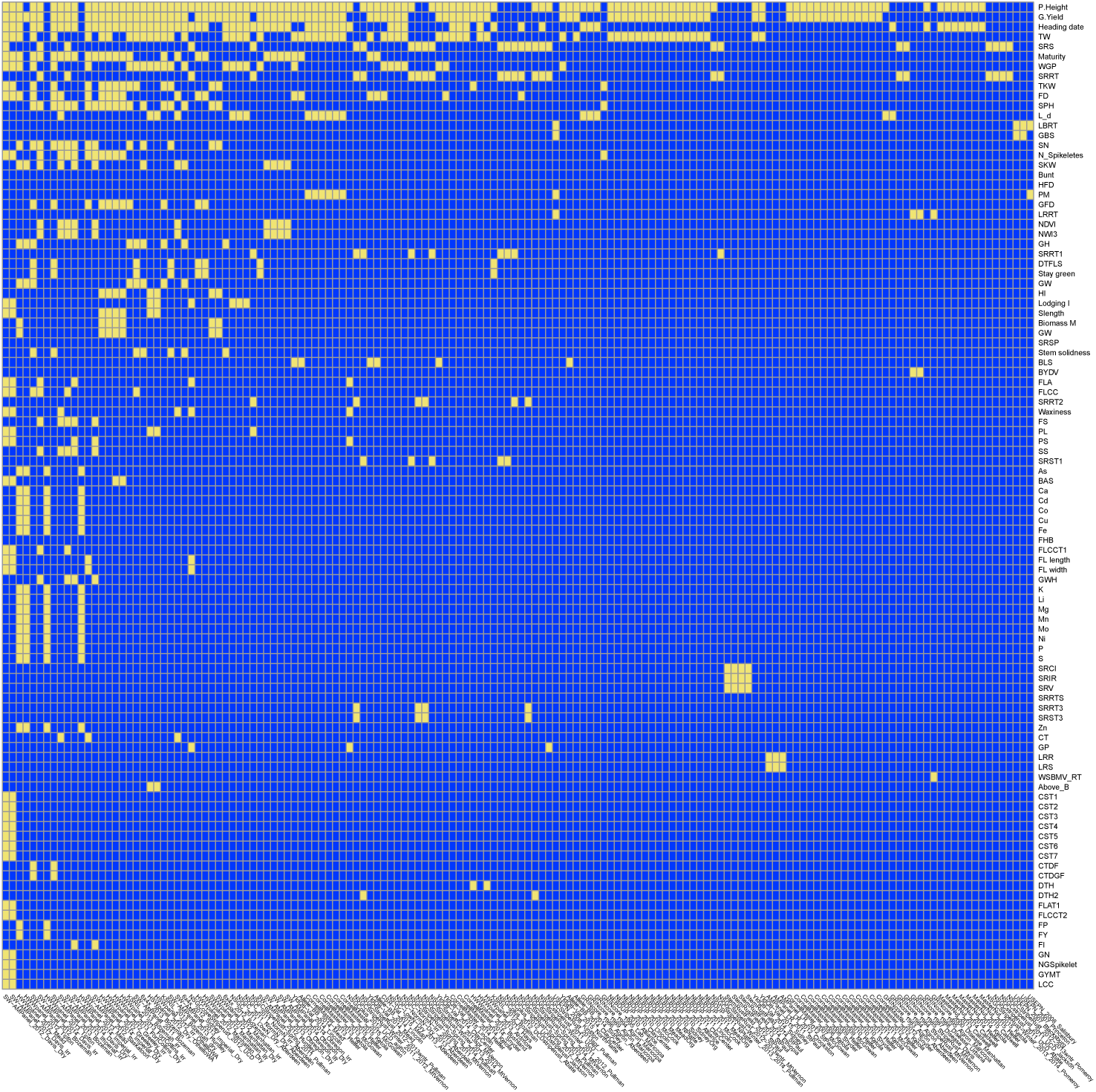
Application 4: Availability of data in 144 phenotypic trials and 95 traits at Triticeae Toolbox for wheat. Yellow shows available data, blue shows unavailable data. The traits and trials are sorted based on availability. Plant height was the most commonly observed trait followed by grain yield.

The correlation matrix for the traits in each trial was calculated and then combined using the Wishart EM-Algorithm. The resulting covariance matrix was used in learning a directed acyclic graph (DAG) using the qgraph R Package [Epskamp et al., 2012].

A more advanced application that involved combining the phenotypic correlation matrices from oat (78 correlation matrices), barley (143 correlation matrices) and wheat (144 correlation matrices) datasets downloaded and selected in a similar way as above were combined to obtain the DAG involving 196 traits in the Supplementary (Supplementary Application 6.1).

## 3 Results

### Application 1- When imputation is not an option: Anchoring independent pedigree-based relationship matrices using a genotypic relation matrix - Potato Data

Figure 6 shows the correlation correlation and mean squared error (MSE) results as either of the sizes of the pedigree matrices and the number of genotypes in the genomic relationship matrices increases. The MSE results for these experiments ranged from 0.004 to 0.017 with a mean of 0.009, and the correlation values ranged from 0.52 to 0.94 with a mean of 0.78.

**Fig. 6.**
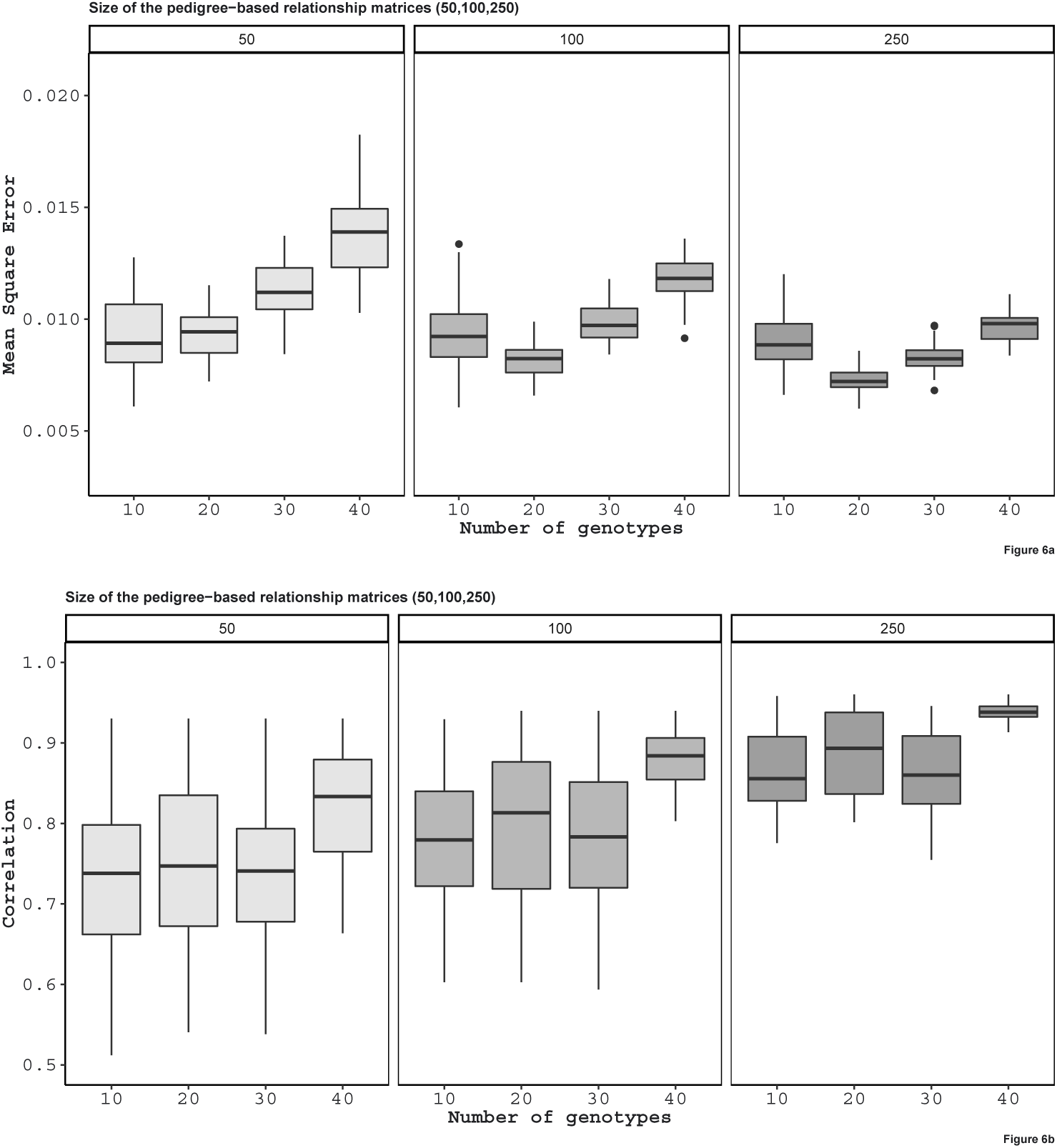
Application 1: For the purpose of this application, the pedigree was split into two pieces although there is only one pedigree. The number of top of the figure is the number of genotypes in each pedigree. Here, we do not know the relationship between one of the pedigree to the another. To learn the relationship between the two, we take 10, 20, 30 and 40 individuals from each group and genotype them by next generation sequencing. The mean square errors and correlation values are the comparison between the two non-overlapping pedigree-based relationship matrices from each sample size, i.e 100 individuals from 50 pedigree based one, and the combined relationship matrix that had 10, 20, 30 and 40 genotypes in each of the pedigrees.

**Fig. 7.**
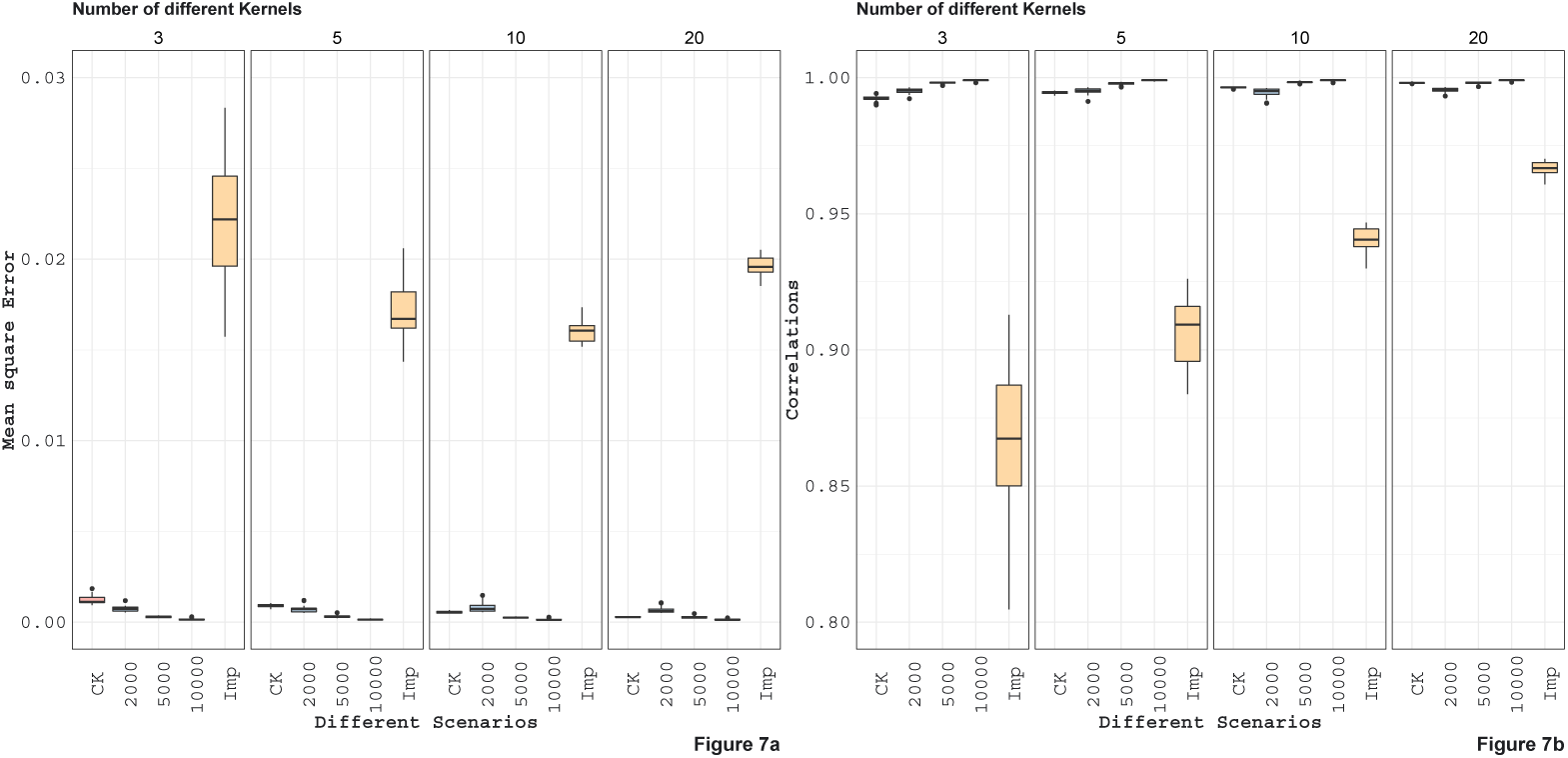
Application 2: Here, we compare marker imputation with our combining relationships matrices approach. Mean square errors and correlations values between the estimated and full genomic relationship matrices are displayed in the boxplots above. The combined relationship matrix (CK) predicts the structure of the population more accurately than the relationship matrix obtained by imputing the genomic features. In addition, when we compare the combined relationship matrix obtained from partially overlapping marker data sets to the relationship matrices obtained from data with a fixed number of markers (2000, 5000, 10000) observed on all individuals we see that combined kernel can be more accurate when the number of partially overlapping marker data sets is large.

### Application 2 - Rice dataset. Combining independent low density marker datasets

The MSE and correlation results for this experiment are given in Figure 7. In general, as the number of independent datasets increases the accuracy of the all of the methods/scenarios increases (decreasing MSEs and increasing correlations). In general, the accuracy of the Wishart EM-algorithm in terms of MSEs ranged from 0.0003 to 0.028 with a mean value of 0.0007. The accuracies measured in correlation ranged from 0.989 to 0.998 with a mean value of 0.995. For the imputation based method MSEs ranged from 0.014 to 0.028 (mean 0.019) and the correlations ranged from 0.805 to 0.970 (mean 0.920).

Figure 8 displays the scatter plot of full genomic relationship matrix (obtained using all 387161 markers) against the one obtained by combining a sample of partial relationship matrices (200 randomly selected genotypes and 2000 randomly selected markers each) over varying numbers of samples (3, 5, 10, 20, 40, and 80 partial relationship matrices). Observed parts (observed-diaginal and observed non-diagonal) of the genomic relationship matrix can be predicted with high accuracy and no bias. As the sample size increase, the estimates get closer to the one obtained using all of the data. We observe that the estimates of the unobserved parts of the relationship are biased towards zero but his bias quickly decreases as the sample size increases.

**Fig. 8.**
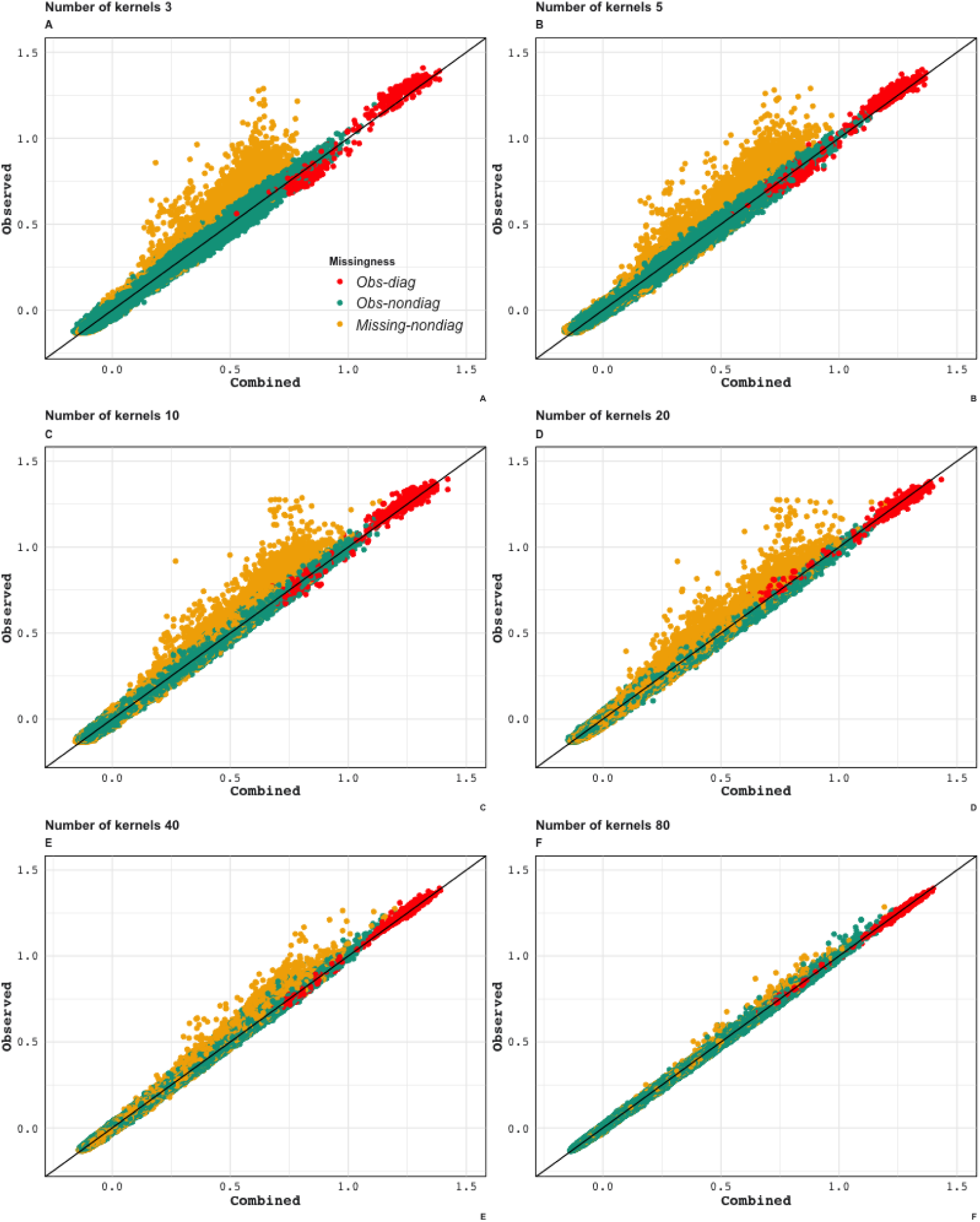
Application 2: Scatter plot of the lower triangular elements of the combined kernel against the kernel calculated from all available markers (Observed). As the number of incomplete datasets increases, both observed and unobserved parts of the relationship can be estimated more precisely. Yellow dots: Genotype relationships that are inferred (not observed in any of the partial relationship matrices that are being combined). Red dots: Diagonal elements of the genotypic relationship matrix. Green dots: Genotype relationships that were observed in one or more of the partial relationship matrices.

### Application 3 - Wheat Data at Triticeae Toolbox. Combining genomic datasets to use in genomic prediction

The results summarized by Figure 9 indicate that when a random sample of genotypes are selected for the test population, the accuracy of the genomic predictions using the combined genomic relationship matrix can be high (Cross-validation scenario 1). Average accuracy for estimating plant height was about 0.68, and for yield 0.58. Lowest accuracy values were for test weight with a mean value of 0.48.. The performance decreases significantly across population predictions (Cross-validation scenario 2). Certain populations were harder to predict, for example, d5, d6, d9. On the other hand, some populations were easier to predict, for instance, from d12 to d16. Average accuracy for estimating plant height was about 0.30, for yield 0.28.

**Fig. 9.**
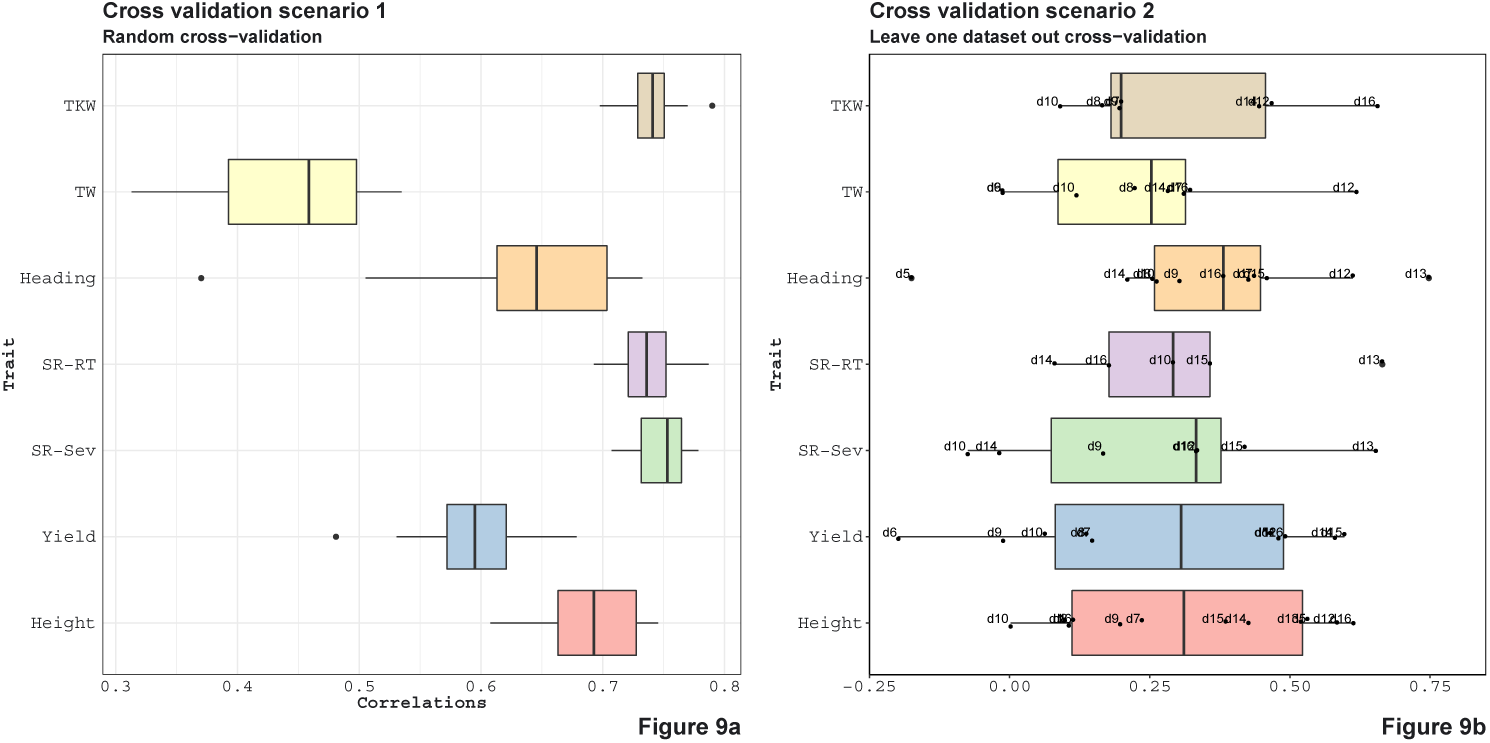
Application 3: Cross-validation scenario 1 is showed in a. For each trait, the available genotypes were split into 10 random folds. The GEBVs for each fold were estimated from a mixed model (See Supplementary Section 5.4 for a description of this model) that was trained on the phenotypes available for the remaining genotypes. Cross-validation scenario 2 is showed in b. Genotypes in each genotypic data are the test and the remaining genotypes are training. In this case, each data that was predicted was also marked on the boxplots. For instance, for plant height, we can predict the phenotypes for the genotypes in d16 with high accuracy when we use the phenotypes of the remaining genotypes as training dataset; on the other hand, we have about zero accuracy when we try to estimate the phenotypes for the genotypes in d10. The accuracy of the predictions under both scenarios were evaluated by calculating the correlations between the GEBVs and the observed trait values.

### Application 4 - Wheat Data at Triticeae Toolbox-Combining Phenotypic Experiments

In this application, we combined correlation matrices obtained from independent phenotypic experiments. Figures 10 and S3 displayed the correlation matrix for the traits in a directed acyclic graph (DAG) and in a heatmap, respectively. In Figure 10 each node represents a trait and each edge represents a correlation between two traits. One of the strength on this representation, is that you can elucidate the correlation between traits that you did not measured in your experiment. For example, among all the traits, grain width (GW) and above ground biomass (Above bm) are positive correlated (blue arrows) with grain yield. In turn, GW is highly positive correlated with biomass at maturity (Biomass M) but negative correlated with harvest index (HI). Negative correlations (red) can also be observed among traits. Traditional inverse correlations such as protein (WGP) and GW can be also observed.

**Fig. 10.**
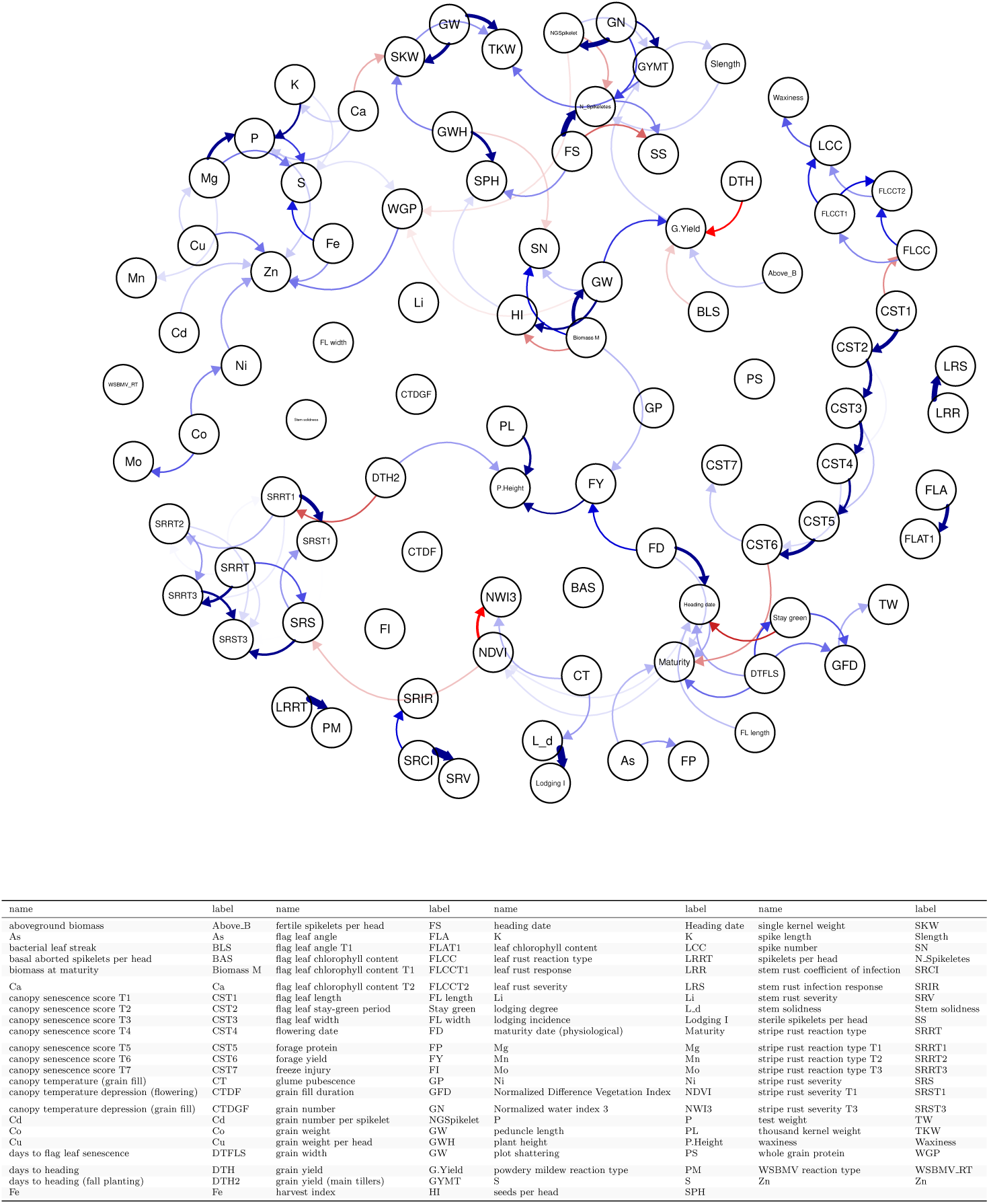
Application 4: Combining the phenotypic correlation matrices from 144 wheat datasets covering 95 traits and illustrating the relationships between traits using the directed acyclic graph as a tool to explore the underlying relationships. Each node represents a trait and each edge represents a correlation between two traits. Blue edges indicate positive correlations, red edges indicate negative correlations, and the width and color of the edges correspond to the absolute value of the correlations: the higher the correlation, the thicker and more saturated is the edge.

Combining datasets by correlation matrices also help to group traits. Figure S3 shows two groups of positively traits. The traits in these two groups are positively correlated within the group but negatively correlated with traits in other groups. For instance, we see that yield related traits such as grain yield, grain weight or harvest index, are positively correlated. On the other hand these traits are negatively correlated with disease related traits such as bacterial leaf streak, stripe rust traits and also with quality traits such as protein and nutrient content.

## 4 Discussion and conclusions

Genomic data are now relatively inexpensive to collect and phenotypes remain to be the primary way to define organisms [Lehner, 2013]. Many genotyping technologies exist and these technologies evolve which leads to heterogeneity of genomic data across independent experiments [Masseroli et al., 2016, Townend, 2018, Lüth et al., 2018]. Similarly, phenotypic experiments, due to the high relative cost of phenotyping, usually can focus only on a set of key traits of interest. Therefore, when looking over several phenotypic datasets, the usual case is that these datasets are extremely heterogeneous and incomplete, and the data from these experiments accumulate in databases [Maiella et al., 2018, Alaux et al., 2018].

This presents a challenge but also an opportunity to make the most of genomic/phenotypic data in the future. In the long term, such databases of genotypic and phenotypic information will be invaluable to scientists as they seek to understand complex biological organisms. Issues and opportunities are beginning to emerge, like the promise of gathering phenotypical knowledge from totally independent datasets for meta-analyses.

To address the challenges of genomic and phenotypic data integration [Suravajhala et al., 2016, Stark et al., 2019], we developed a simple and efficient approach for integrating data from multiple sources. This method can be used to combine information from multiple experiments across all levels of the biological hierarchy such as microarray, gene expression, microfluidics, and proteomics will help scientists to discover new information and to develop new approaches.

For example, Figure 7 shows that we can estimate the full genomic relationship matrix more precisely from 10 independent partially overlapping datasets of 200 genotypes and 2000 markers each than estimating from a dataset (for the combined set of genotypes) that has 2000 fixed markers. Twenty independent genomic datasets of 200 genotypes and 2000 markers is as good as one genomic dataset with 5000 markers. When we compare it to the rest of the entries, imputation is the least effective for estimating the unobserved parts of the genomic relationship matrix. This suggests that accounting for incomplete genetic relationships would be a more promising approach than estimating the genomic features by imputation and then calculating the genomic relationship matrix.

Figure 6 shows we can accurately estimate the unobserved relationships among the genotypes in two independent pedigree based relationship matrices by genotyping a small proportion of the genotypes in these datasets. For instance, the mean correlation for the worst case setting (50 genotypes in each pedigree and 10 from each of the pedigree genotyped) was 0.72. This value increased all the way up to 0.94 for the best case (250 genotypes in each pedigree and 40 from each of the pedigree genotyped).

The selection in GS is based on GEBVs and a common approach to obtaining them involves the use of a linear mixed model with a marker-based additive relationship matrix. If the phenotypic information corresponding to the genotypes in one or more of the component matrices are missing then the genotypic value estimates can be obtained using the available phenotypic information. In this sense, the combined genomic information links all the genotypes and the experiments.

Imputation has been the preferred method when dealing with incomplete and datasets [Browning, 2008, Browning and Browning, 2009, Howie et al., 2011, Druet et al., 2014, Erbe et al., 2016]. However, imputation can be inaccurate if the data is very heterogeneous [Van Buuren et al., 2011]. In these cases, as seen in applications above, the proposed approach which uses the relationships instead of the actual features seems to outperform imputation for inferring genomic relationships. Besides, the methods introduced in this article are useful even when imputation is not feasible. For example, two partially overlapping relationship matrices, one pedigree-based and the other can be combined to make inferences about the genetic similarities of genotypes in both of these datasets (Figure 6).

There are also limitations to our approach. In particular, when we combine data using relationship matrices original features (markers) are not imputed. Our method may not be the best option when inferences about genomic features are needed, such as in GWAS. We can address this issue by imputing the missing features using the combined relationship matrix, for instance, using a k-nearest neighbor imputation [Hastie et al., 2001] or by kernel smoothing. Moreover, if the marker data in the independent genomic studies can be mapped to local genomic regions, then the combined relationship matrices can be obtained for these genomic regions separately. Then a kernel based model such as the ones in Yang et al. [2008], Akdemir and Jannink [2015] can be used for association testing. The nature of missingness in data will affect our algorithms performance. Inference based on approaches that ignore the missing data mechanisms are valid for missing completely at random, missing at random but probably not for not missing at random [Little and Rubin, 2002, Rubin, 1976].

### 4.1 Software and data availability

The software was written using C++ and R and an R (R Core Team [2019]) package **CovCombR** [Akdemir et al., 2020] is made available publicly. The code and data for replicating some of the analysis can be requested from the corresponding author.

## Supplementary Materials

## 5 Supplementary Methods

### 5.1 Wishart EM-Algorithm

The Wishart EM-Algorithm maximizes the likelihood function for a random sample of incomplete observations from a Wishart distribution with fixed degrees of freedom since it is an EM-Algorithm [Dempster et al., 1977, 1981]. To the best of our knowledge, this is the first study that derives the EM-Algorithm for the following case. Let 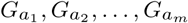 be independent and partial realizations from a Wishart distribution with a known degrees of freedom *ν > n* and covariance parameter *Ψ* = *Σ/ν*. Expectation of each *G*_*a*_ is therefore equal to *Σ*_*a*_.

The likelihood function for the observed data can be written as

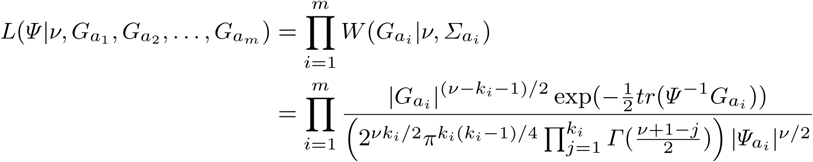

The log-likelihood function with the constant terms combined in *c* is given by

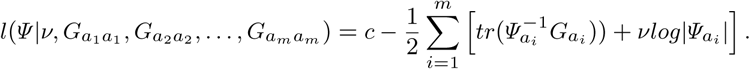

Complementing each of the observed data with the missing data components *G*_*B*_ = (*G*_*ab*_, *G*_*b*_), we can write the log-likelihood for the complete data up to a constant term as follows:

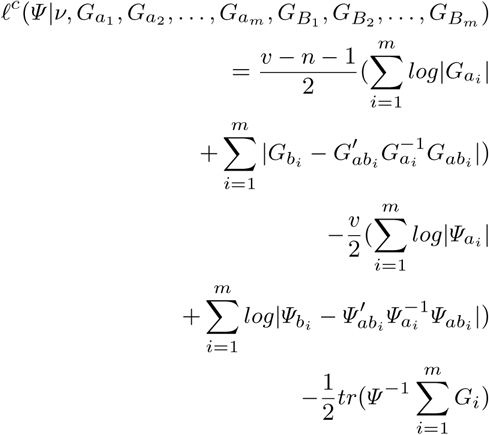

The expectation step of the EM-Algorithm involves calculating the expectation of the complete data log-likelihood conditional on observed data and the value of *Ψ* at iteration *t* which we denote by *Ψ* (*t*).

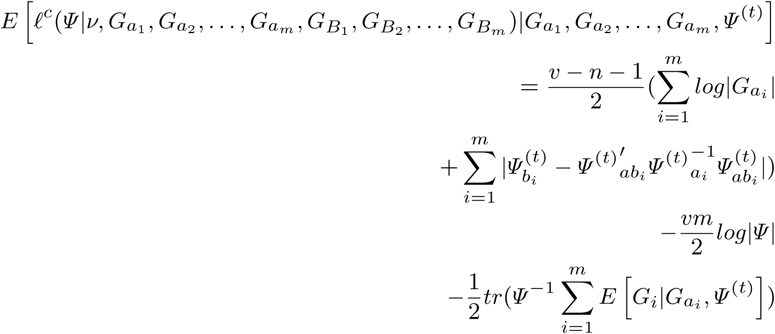

The maximization step of the EM algorithm which updates *Ψ* ^(*t*)^ to *Ψ* ^(*t*+1)^ by finding *Ψ* that maximizes the expected complete data log-likelihood. (Using [Anderson, 1984b, Lemma 3.3.2]) The solution is given by:

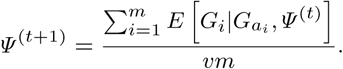

We need to calculate 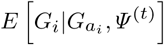 for each *i. G* is partitioned as

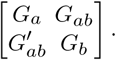

We assume a similar partitioning for *Ψ*.

Firstly, 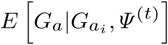is *G*_*a*_. Secondly, 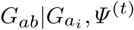 has a matrix-variate normal distribution with mean 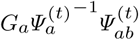 (the covariance of the vectorized form is given by 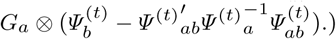

To calculate the expectation of *G*_*b*_, note that we can write this term as 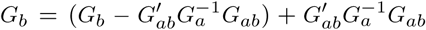. The distribution of the first term is independent of *G*_*a*_ and *G*_*ab*_ and is a Wishart distribution with degrees of freedom *ν* − *n*_*a*_ and covariance parameter 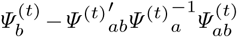. The second term is an inner product 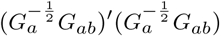. The distribution of 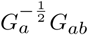 is a matrix-variate normal distribution with mean 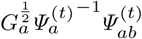 and covariance is given by 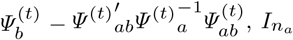 for the columns and rows correspondingly. The expectation of this inner-product is 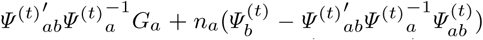. Therefore, the expected value of *G*_*b*_ given 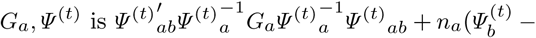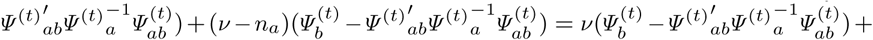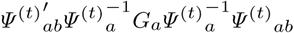 This leads to the update equation:

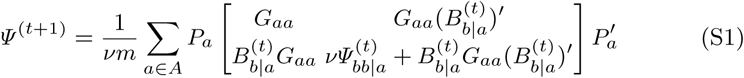

where 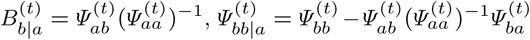, *a* is the set of genotypes in the given partial genomic relationship matrix and *b* is the set difference of *K* and *a*. The matrices *Pa* are permutation matrices that put each matrix in the sum in the same order. The initial value, *Σ*^(0)^ is usually assumed to be an identity matrix of dimension *n*.

During the steps of the Wishart EM-Algorithm we might encounter a matrix *Ψ* which is not positive definite. There are two strategies to deal with this case: allow *Ψ* to be non definite but replace it with a near positive definite matrix after last iteration, 2) force *Ψ* to be positive definite at each iteration by replacing it with a near positive definite matrix. We have used the second approach in our implementations.

#### Asymptotic standard errors

Once the maximizer of 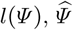, has been found, the asymptotic standard errors can be calculated from the information matrix of *Ψ* evaluated at 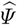. The log-likelihood is given by:

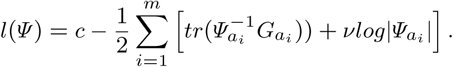

First derivative with respect to the *jk*th element of *Ψ* is given by

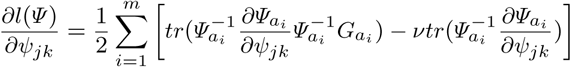

The derivative of the above with respect to the *lh*th element of *Ψ* is given by

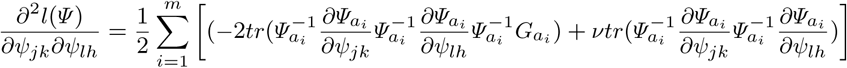

Expected value of the second derivative is given by

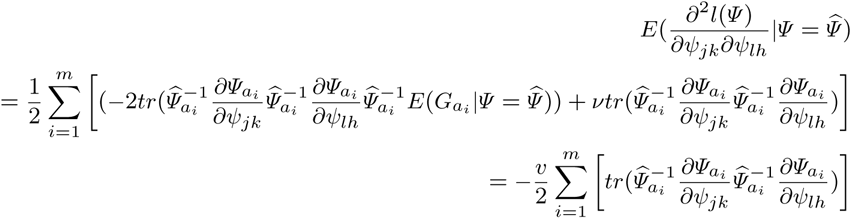

Therefore, the information matrix is given by

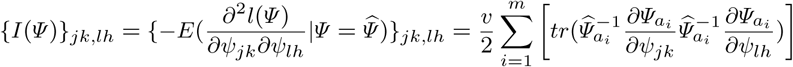

### 5.2 Some Properties of Matrix Normal and Wishart Distribution

The following results and their derivations are given in classic multivariate statistics textbooks such as [Anderson, 1984a] and [Gupta and Nagar, 2000, Kollo and von Rosen, 2006] and are used in the derivation of the Wishart EM-Algorithm.

**–** [Kollo and von Rosen, 2006, Theorem 2.2.9] Let *X ∼N*_*p,n*_(*M, Σ, Ψ*). Then, *E*[*XAX′*] = *tr*(*ΨA*)*Σ* + *MAM ′*.
**–** [Kollo and von Rosen, 2006, Theorem 2.4.12.]Let *G ∼ W*_*n*_(*ν, Ψ*) with *Ψ* and *ν > n*.
**–**Density

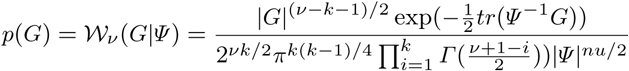
**–***E*(*G*) = *νΨ*
**–***G*_1|2_ is independent of (*G*_12_, *G*_22_);
**–***G*_22_ *∼ W*_*q*_ (*ν, Ψ*_22_
**–**The conditional distribution of *G*_12_ given *G*_22_ is multivariate Gaussian 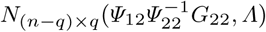 where 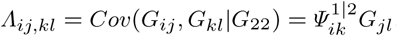.

### 5.3 Genomic features, distances and kernel matrices

Let *M* be the *n × m* matrix of biallelic marker allele dosages for *n* genotypes and *m* markers, and let *n < m*. The vector of estimates of allele probabilities is given by 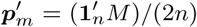. Let 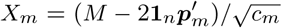be the feature matrix where 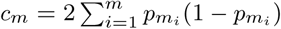. An additive relationship matrix can be written 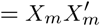 [VanRaden, 2008]. This matrix is singular.

A similar relationship matrix that is nonsingular can be obtained by changing the centering and scaling of the allele dosages matrix. Let 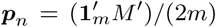. Let 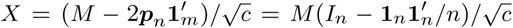 be the feature matrix where 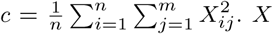. *X* is the row centered feature matrix scaled by the mean square root of total average heterozygosity for the genotypes. We also use the notation *G*_*A*_(*X*) = *XX′* and note that *G*_*A*_(*X*) can be calculated from by covariance matrix for the genotypes of the marker allele dosages matrix *M* by dividing it by the mean of its diagonal elements (abusing notation, this can be expressed as *G*_*A*_(*X*) = *cov*(*M ′*)*/mean*(*diag*(*cov*(*M ′*))).). This matrix is non-singular whenever the number of independent features in the data are larger than the sample size. The mean of the diagonals of this relationship matrix is one. More importantly, the same formulation applies to all types of genomic features. For instance, we can use the same formulation for marker data with higher ploidy levels, or with other forms of genomic data such as the expression data.

For each pair of genotypes ((*i, j*) : *i, j ∈*(1, 2, …, *n*)) in *M*, the squared Euclidean distance using the corresponding a feature matrix *X* = (***x***_1_, ***x***_2_, …, ***x***_*n*_)*′* can be written as

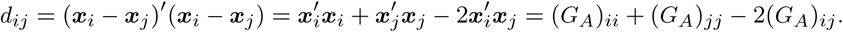

The squared distance matrix is defined by *D*(*X*) = (*d*_*ij*_) and can be calculated from the additive relationship matrix *G*_*A*_(*X*) = *XX′* as

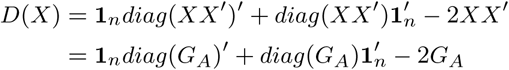

Moreover, since **1***′X* = **0** and 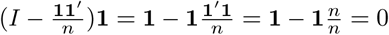, we have

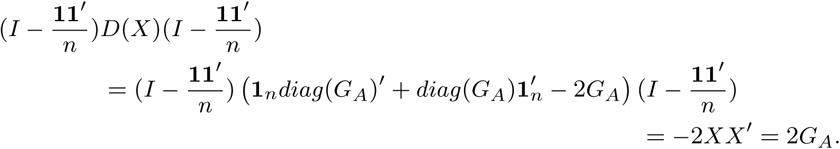

Therefore, given *D*(*X*) and letting 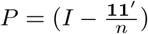 the additive relationship matrix can also be calculated by

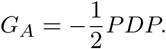

The genomic relationship matrices need not be additive. RKHS regression extends additive relationship based SPMMs by allowing a wide variety of kernel matrices, not necessarily additive in the input variables, calculated using a variety of kernel functions. A kernel function, *k*(., .) maps a pair of input points ***x*** and ***x****′* into real numbers. It is by definition symmetric (*k*(***x, x****′*) = *k*(***x****′*, ***x***)) and non-negative. Given the inputs for the *n* genotypes we can compute a kernel matrix *G* whose entries are *G*_*ij*_ = *k*(***x***_*i*_, ***x***_*j*_). The linear kernel function is given by *k*(***x***; ***y***) = ***x****′****y***. The polynomial kernel function is given by *k*(***x***; ***y***) = (***x****′****y*** + *c*)^*d*^ for *c* and *d ∈ R*. Finally, the Gaussian kernel function is given by *k*(***x***; ***y***) = *exp*(−*h*(***x****′* − ***y***)*′*(***x****′* − ***y***)) where *h >* 0. The common choices for kernel functions are the linear, polynomial, Gaussian kernel functions, though many other options are available [Schölkopf and Smola, 2005, Endelman, 2011].

The relationship between the Euclidean distance matrix and the corresponding Gaussian kernel is given by

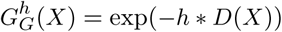

and

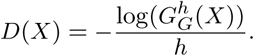

An important advantage of using similarity or distance matrices over the original features is that similarity of distance matrices can be calculated for variables of different type (categorical, rank, or interval-scale data). The relationship of the feature matrix, and the additive kernel and Euclidean distance allows us to generalize the additive relationship matrix to general genomic data (not necessarily marker allele dosages).

### 5.4 Mixed models and genomic relationship matrices

Let’s start by describing how we can use a single combined genomic data. The discussion below will be biased towards a discussion variance components / mixed modeling approach since this has a special place in quantitative genetics. Mixed models have been used as a formal way of partitioning the variability observed in traits into heritable and environmental components, it is also useful in controlling for population structure and relatedness for genome-wide association studies (GWAS). However, some of the methods that are proposed can be used in other forms of statistical analysis, for instance, for descriptive purposes or in general statistical learning.

In a mixed model, genetic information in the form of a pedigree or marker allele frequencies can be used in the form of an additive genetic similarity matrix that describes the similarity based on additive genetic effects (GBLUP). For the *n ×* 1 response vector ***y***, the GBLUP model can be expressed as

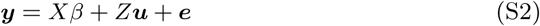

where *X* is the *n × p* design matrix for the fixed effects, *β* is a *p ×* 1 vector of fixed effect coefficients, *Z* is the *n × q* design matrix for the random effects; the vector random effects (***u****′*, ***e****′*)*′* is assumed to follow a multivariate normal (MVN) distribution with mean **0** and covariance

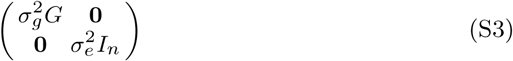

where *G* is the *q × q* additive genetic similarity matrix. In this model, the labels of the genotypes (that are listed in the rows and columns of the relationship matrix *G*) define a factor variable with levels equal to the labels. The matrix *Z* is the design matrix that links the observed response in the experiment to these levels. The model (S2) is equivalent to a MM in which the additive marker effects are estimated via the following model (rr-BLUP):

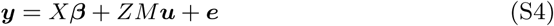

where *X* is the *n × p* design matrix for the fixed effects, *β* is a *p ×* 1 vector of fixed effect coefficients, *Z* is the *n × q* design matrix for the random effects *M* is *q × m* marker allele frequency centered incidence matrix; (***u****′*, ***e****′*)*′* follows a MVN distribution with mean **0** and covariance

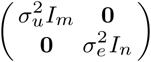

Note that the scale of the genomic relationship matrix is irrelevant for genomic prediction or for family structure correction in mixed model-based association studies. However, this quantity is important for the calculation of narrow-sense heritability. In this case, setting the average of the diagonals of the relationship makes it, in a way, compatible with the broad sense heritability calculations based on an identity relationship matrix for genotypes that already has a mean of its diagonal elements equal to one. In addition, the standard formulations of the marker-based additive matrix models used in the literature can be generalized to incorporate more complex genetic and environmental covariates.

## 6 Supplementary Applications

### 6.1 Experiments with simulated data

#### Supplemenatry Application 1- Simulation study: Inferring the combined covariance matrix from its parts

To establish that a combined relationship can be inferred from realizations of its parts, we have conducted the following simulation study: In each round of the simulation, the true parameter value of the genomic relationship matrix was generated as 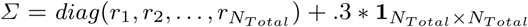 where *r*_*i*_ were independently generated as 1 + .7 *\* u*_*i*_ with *u*_*i*_ a realization from the uniform distribution over (0, 1). *Σ* was then adjusted by dividing it with the mean value of its diagonal elements. This parameter was taken as the covariance parameter of a Wishart distribution with degrees of freedom 300 and *N*_*kernel*_ samples from this distribution are generated. After that, each of the realized relationship matrices was made partial by leaving a random sample of 10 to 40 (this number was also selected from the discrete uniform distribution for integers 10 to 40) genotypes in it. These partial kernel matrices were combined using the Wishart EM-Algorithm iterated for 50 rounds (each round cycles through the partial relationship matrices in random order). The resultant combined relationship matrix 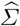 was compared with the corresponding parts of the parameter *Σ*^3^ by calculating the mean squared error between the upper diagonal elements of these matrices. This experiment was replicated 10 times for each value of *N*_*Total*_ *∈* {40, 80, 150, 300} and *N*_*kernel*_ *∈* {40, 80, 150, 300}.

The results of this simulation study are summarized in Figure S1. For each covariance size, the MSE’s decreased as the number of incomplete samples increased. On the other hand, as the size of the covariance matrix increased the MSEs increased.

#### Supplementary Application 2-Simulation study: Likelihood Convergence

The Wishart EM-Algorithm maximizes the likelihood function for a random sample of incomplete observations from a Wishart distribution. The derivation of this feature is given in the Supplementary. In this application, we explore the convergence of the algorithm for several instances starting from several different initial estimates.

The application is composed of 10 experiments each of which starts with a slightly different assumed Wishart covariance parameter^4^. For each true assumed covariance matrix, we have generated 10 partial samples including between *n*_*min*_ and *n*_*max*_ genotypes (random at discrete uniform from *n*_*min*_ to *n*_*max*_) each using the Wishart distribution. *n*, the total number of genotypes in the assumed relationship matrix was taken to be 100 or 1000. Corresponding to this two matrix sizes the *nmin* and *nmax* are taken as 10 and 25 or 100 and 250. These 10 matrices are combined using the Wishart EM-Algorithm 10 different times each times using a slightly different initial estimate of the covariance parameter^5^. We record the path of the log-likelihood function for all these applications.

**Fig. S1.**
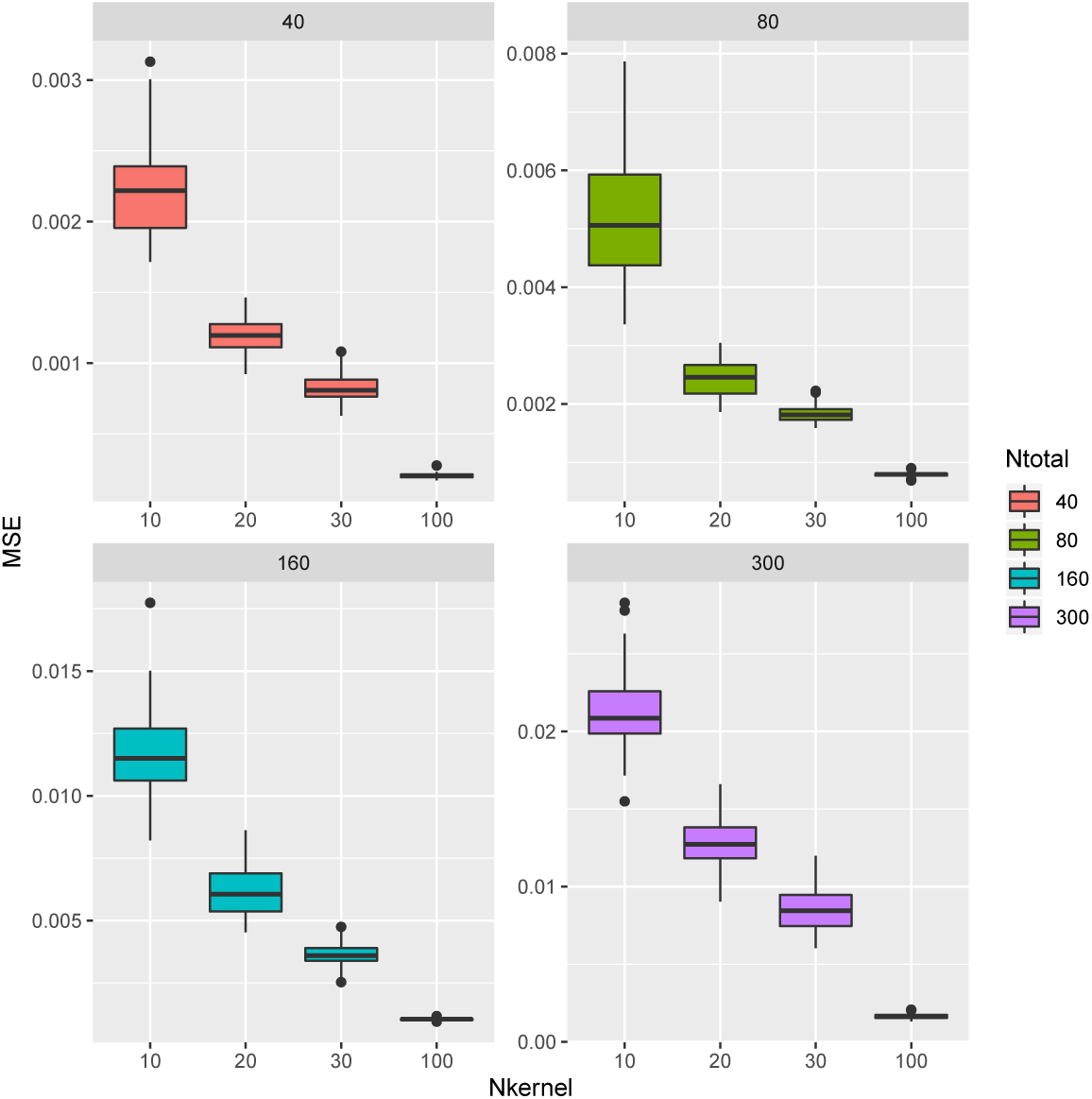
Application 1 - MSE’s for estimating correlation parameters based on partial samples for *N*_*T otal*_ *∈* {40, 80, 150, 300} (number of variables in the covariance matrix) and *N*_*kernel*_ *∈{*40, 80, 150, 300} (number of incomplete covariance matrix samples). Each incomplete covariance matrix was had a random size between 10 to 40. The MSE’s are calculated over 10 replications of the experiment.

At each instance of the parameter and a particular sample, the likelihood functions converged to the same point (See Figure S2). We have not observed any abnormalities in convergence according to these graphs.

#### Heatmap for 95 wheat traits

**Phenotypic network for 186 traits based on phenotypic correlations (Wheat, Barley, and Oat Phenotypic Trials from Triticeae Toolbox)**

### 6.2 Supplementary Figures

**Fig. S2.**
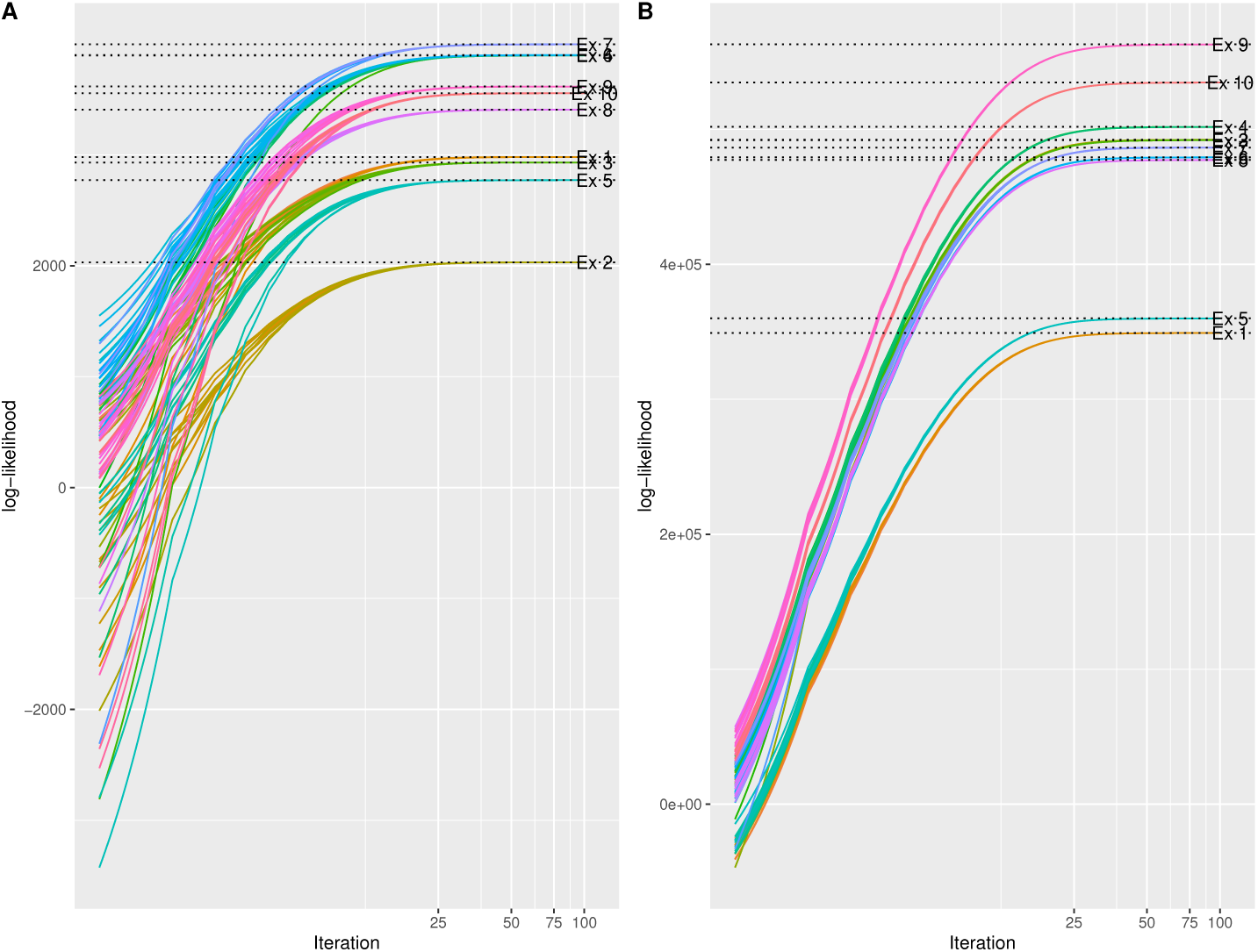
Application 2 - Convergence of log-likelihood function: Each color represents a different experiment. In each experiment, a sample of incomplete covariance matrices from a Wishart distribution were combined using the Wishart EM-Algorithm starting from 10 different slightly different random initial estimates. *n*, the total number of genotypes in the assumed relationship matrix was taken to be 100 (**A**) or 1000 (**B**).

**Fig. S3.**
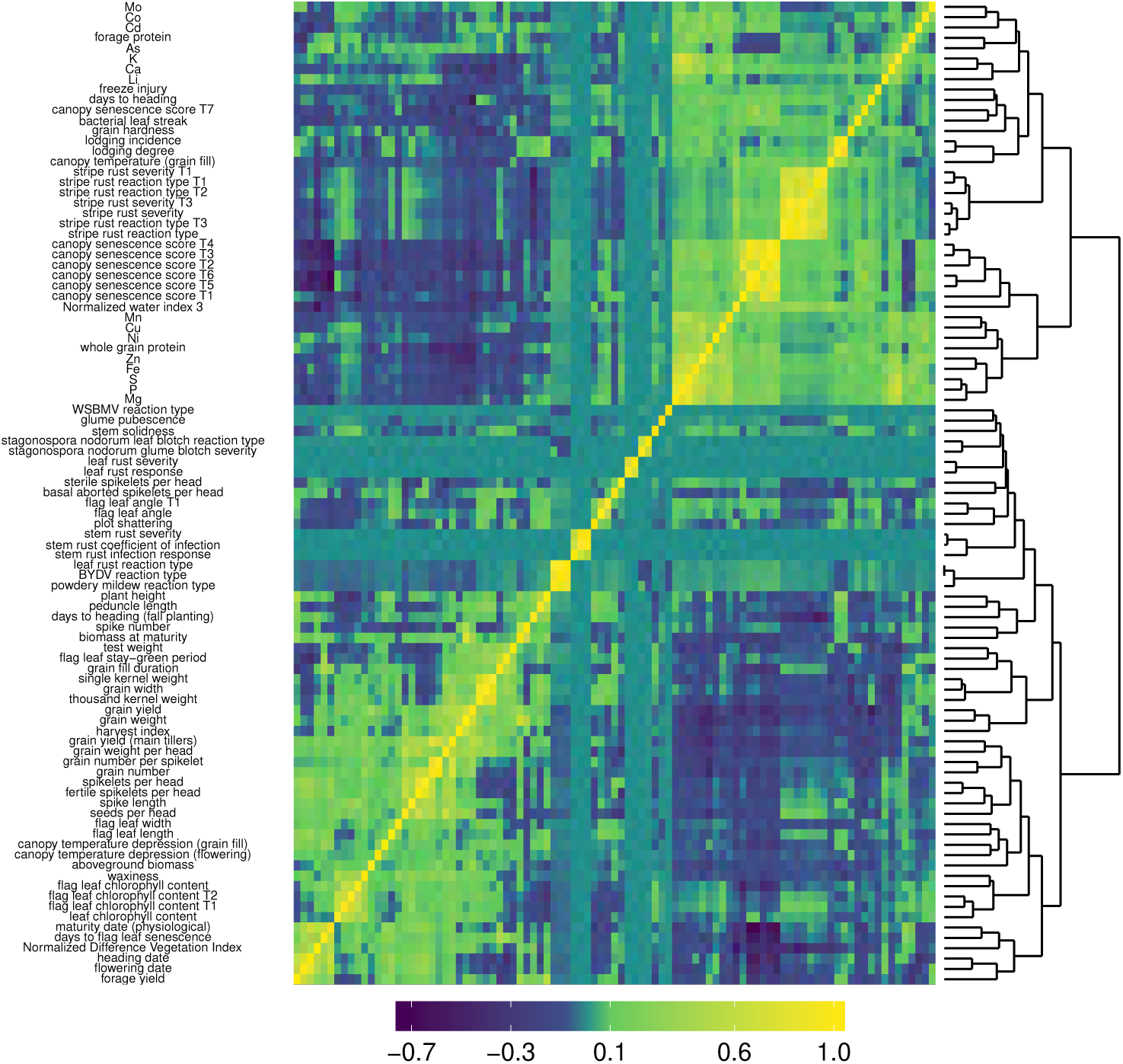
Triticeae dataset: Combining the phenotypic correlation matrices from 144 wheat datasets covering 95 traits. Clustered heatmap of Pearson correlation coefficients provides a global overview of phenotypic correlation across wheat traits. Yellow denotes high correlation, dark green high anti-correlation.

**Fig. S4.**
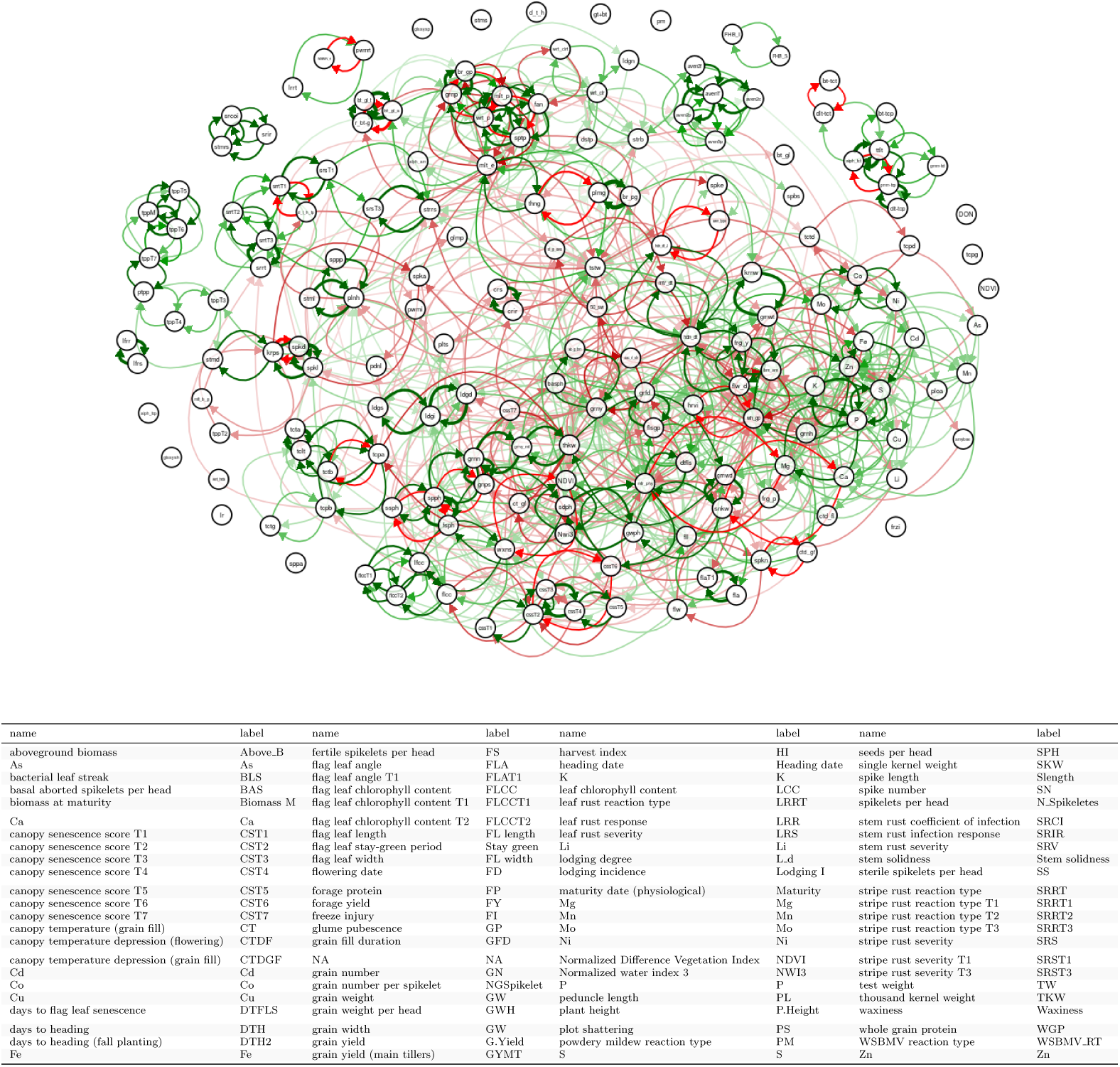
Triticeae datasets: Combining the phenotypic correlation matrices from oat (78 correlation matrices), barley (143 correlation matrices) and wheat (144 matrices) datasets downloaded and selected in a similar way as in Application 4 were combined to obtain the DAG involving 196 traits.

**Fig. S5.**
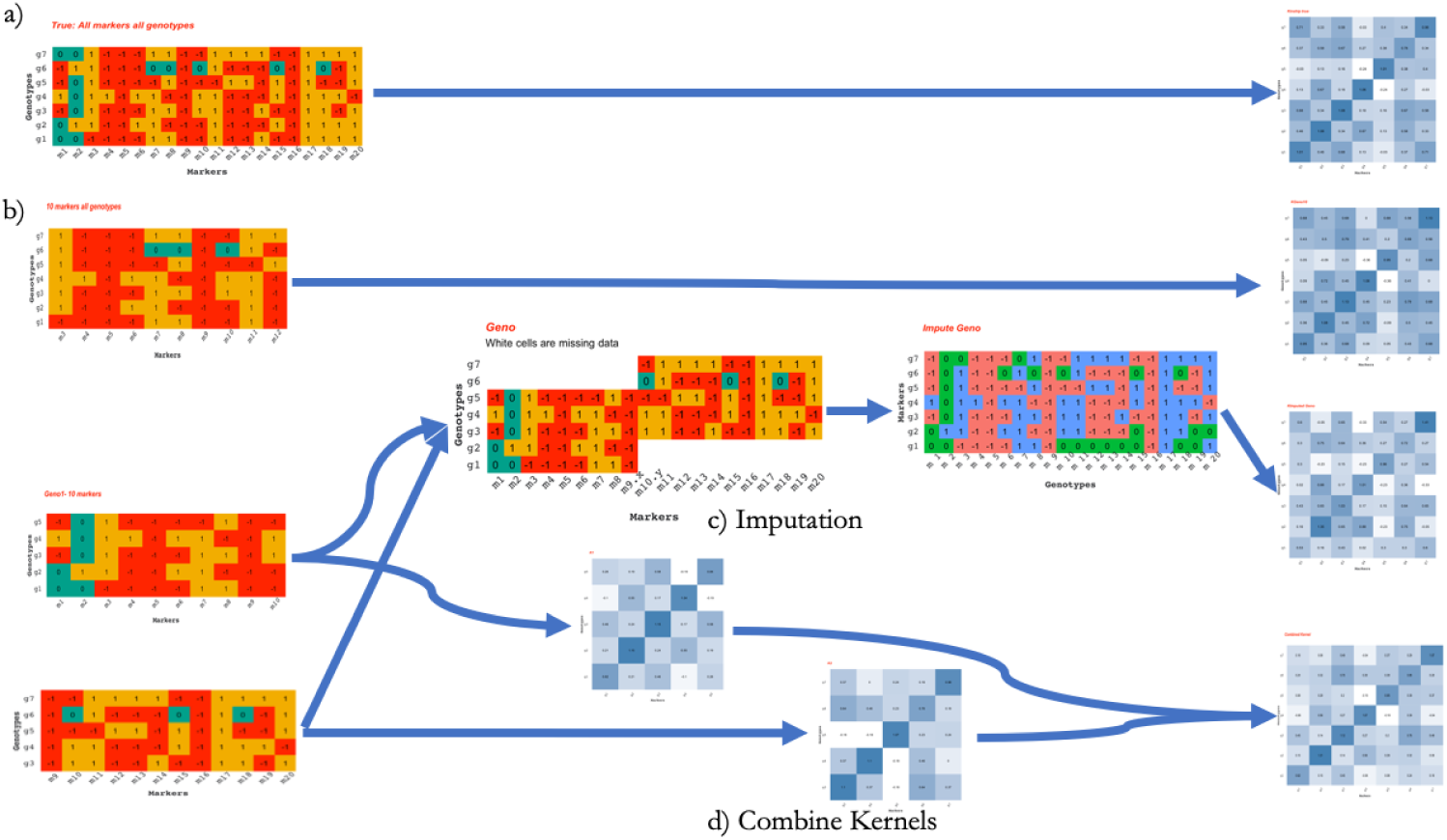
Pictorial representation for some of the different scenarios in Application 2 with reduced number of markers, genotypes and number of independent marker datasets. a) Assumed truth, b) All genotpes using 10 markers, c) Imputation of 2 independent marker datasets, d) Combining the relationship matrices from 2 independent marker datasets.

**Fig. S6.**
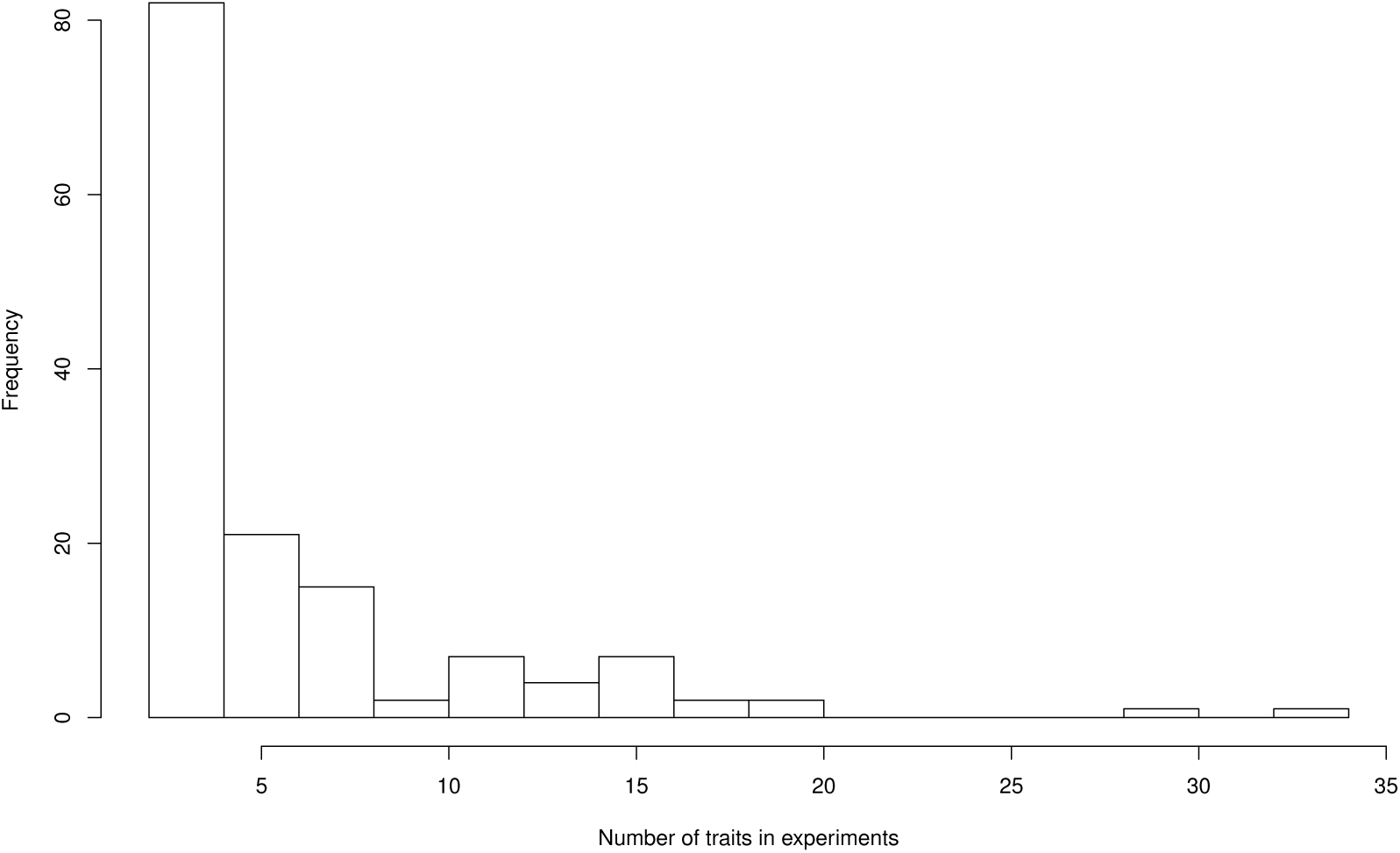
Triticeae dataset: The distribution of the numbers of traits in 144 phenotypic trials at Triticeae Toolbox for wheat. The mean and the median of the number of traits in these trials were 5.9 and 4 correspondingly.

**Fig. S7.**
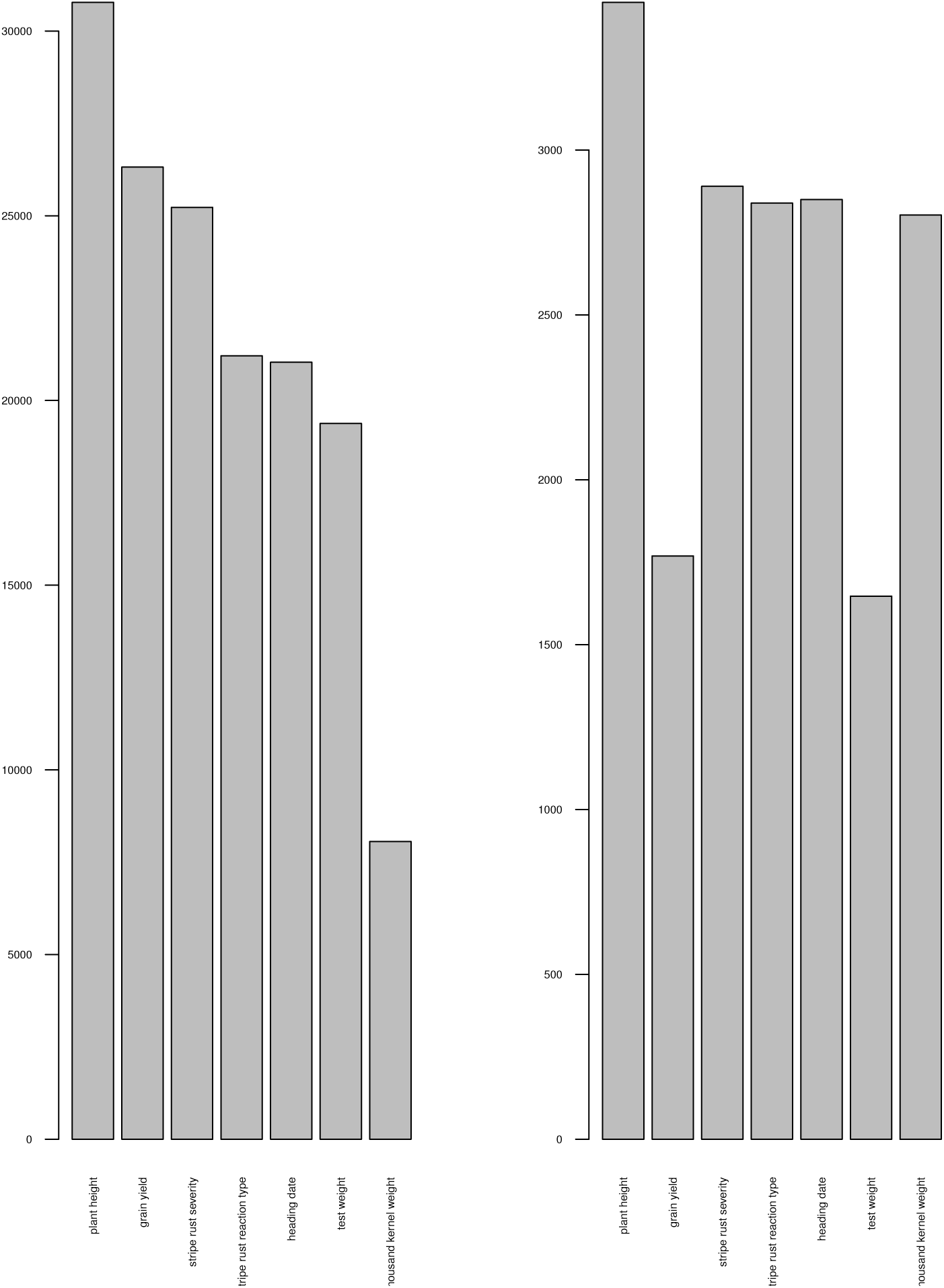
Triticeae dataset: Number of phenotypic observations (left) and the number of genotypes available in Triticeae Toolbox for a set of 7 selected traits for the 9102 genotyes in the combined relationship matrix.

## Footnotes

1 In what follows, we will refer to genetic relationship matrices that measure how genotypes are related (See Supplementary Section 5.3 for a description of how to calculate a genetic relationship matrix from genome-wide markers (genomic relationship matrix)). However, a theme in this article is that a genetic relationship matrix is a special kind of covariance matrix. Therefore, the same arguments below apply to covariance matrices that measure the relationship between traits or features.

2 We used correlations instead of covariances because the phenotypic experiments were very heterogeneous in terms of the variances of the different traits.

3 In certain instances, the union of the genotypes in the parts did not recover all of the *N*_*T otal*_ genotypes, therefore this calculation was based on the recovered part of the full genomic relationship matrix

4 *Σ* = *diag*(***b*** + 1) + .21_*n×n*_ where ***b***_*i*_ for *i* = 1, 2, …, *n* are i.i.d. uniform between 0 and 1.

5 *Σ*_0_ = *diag*(.5***b*** + 1) + .3 *\*b*_0_1_*n×n*_ where ***b***_*i*_ for *i* = 0, 2, …, *n* are i.i.d. uniform between 0 and 1.

